# OpaR exerts a dynamic control over c-di-GMP homeostasis and *cpsA* expression in *V. parahaemolyticus* through its regulation of ScrC and the trigger phosphodiesterase TpdA

**DOI:** 10.1101/2023.01.10.523516

**Authors:** David Zamorano-Sánchez, Jesús E. Alejandre-Sixtos, Adilene Arredondo-Hernández, Raquel Martínez-Méndez

## Abstract

The second messenger cyclic dimeric guanosine monophosphate (c-di-GMP) plays a central role in controlling decision making processes of vital importance for the environmental survival of the human pathogen *Vibrio parahaemolyticus*. The mechanisms by which c-di-GMP levels are dynamically controlled in *V. parahaemolyticus* are poorly understood. Here we report our findings regarding the involvement of OpaR in controlling c-di-GMP metabolism in planktonic and surface-attached cells through controlling the expression of the trigger phosphodiesterase (PDE) TpdA and other PDEs such as ScrC. Our results revealed that OpaR negatively modulates the expression of *tpdA* by maintaining a baseline level of c-di-GMP. The OpaR-regulated PDEs ScrC, ScrG and VP0117 enable the upregulation of *tpdA*, to a different degree, in the absence of OpaR. We also found that TpdA plays the dominant role in c-di-GMP degradation under planktonic conditions compared to the other OpaR-regulated PDEs. In cells growing over solid media the dominant c-di-GMP degrader role is played by ScrC for 72 hours and passes to TpdA after 96 hours of growth. We also report negative and positive effects of the absence of OpaR on *cpsA* expression in cells growing over solid media or forming biofilms over glass, respectively. These results suggest that OpaR can act as a double-edged sword to control c-di-GMP accumulation and *cpsA* expression positively or negatively in response to poorly understood environmental factors. Finally, through an *in-silico* analysis we point out outlets of the OpaR regulatory module that can impact decision making during the motile to sessile transition in *V. parahaemolyticus*.

**Importance:** The second messenger c-di-GMP is extensively used by bacterial cells to control crucial social adaptations such as biofilm formation. Here we explore the role of the quorum-sensing regulator *OpaR*, from the human pathogen *V. parahaemolyticus*, on the dynamic control of c-di-GMP signaling. We found that OpaR can regulate positively or negatively c-di-GMP accumulation depending on the growth conditions. This dual role has not been reported for orthologues of OpaR, such as HapR from *V. cholerae*. OpaR controls c-di-GMP homeostasis through PDEs that are absent in *V. cholerae*, pointing toward further differences in c-di-GMP signaling in these two pathogens. It is important to investigate the origins and consequences of these differences to better understand pathogenic bacterial behavior and its evolution.

## Introduction

The *Vibrio* genus includes several bacterial aquatic species that continuously threaten human health worldwide. *Vibrio parahaemolyticus* is one of the most common etiological agent for gastroenteritis caused by raw seafood consumption (1). This pathogen also impact human activities such as shrimp farming due to its ability to cause an acute hepatopancreatic necrosis disease in these organisms (2, 3). The ability of this pathogen to adapt to the ever-changing ecosystems where it dwells is of paramount importance for its colonization success and its persistence and dissemination in the environment.

Like for many pathogens, it is likely that the ability of *V. parahaemolyticus* to form complex multicellular associations within a self-produced exopolymeric matrix, known as biofilms, plays a key role in its ability to resist adverse conditions such as antimicrobial insults and nutrient scarcity (4, 5). Our knowledge of the process of biofilm formation and its regulation in the *Vibrio* genus has been mostly obtained from the study of the human pathogen *Vibrio cholerae* (6). Although several of the paradigms of biofilm formation in *V. cholerae* are true for other *Vibrio* species, there are important particularities in mono-species biofilms within the *Vibrio* genus that justify the need to study them in a case to case basis (7). Increasing our understanding of the process of biofilm formation and biofilm development could enable a more educated design of contention strategies to combat *V. parahaemolyticus* outbreaks.

A general element of study within the biofilm formation and regulation field is the role of the second messenger cyclic dimeric guanosine monophosphate (c-di-GMP). This intracellular messenger plays a central role in the process of decision making in bacteria including the choice between a planktonic or a sessile life-style (8, 9). C-di-GMP is synthesized by diguanylate cyclases (DGCs) that have a catalytic GGDEF domain and it is degraded by specific phosphodiesterases (PDEs) with a catalytic EAL or HD-GYP domain (8). C-di-GMP signaling modules are typically composed by DGCs and PDEs and one or multiple effectors that can interact with c-di-GMP and in consequence alter a physiological response (10). Changes in c-di-GMP levels affect the transcriptomic profile of *V. cholerae* and its ability to form biofilms (11, 12). Multiple PDEs and DGCs have been shown to participate in controlling the intracellular pool of c-di-GMP in *V. cholerae;* however, less is known about the cues that result in changes in the activity or abundance of the components of these c-di-GMP signaling modules (11, 13–15). A key modulator of c-di-GMP accumulation and biofilm formation in *V. cholerae* is the quorum sensing master regulator HapR, which negatively controls these processes through its ability to affect the transcription of genes related to c-di-GMP metabolism and exopolysaccharide biosynthesis (13, 16–18). This regulator is conserved in species within the *Vibrio* genus; in *V. parahaemolyticus* this regulator was named OpaR due to its involvement in controlling colony opacity, a phenotype that is influenced by the ability to produce a capsular polysaccharide named CPS (19, 20). Opposite to HapR, which was shown to repress exopolysaccharide biosynthesis and c-di-GMP accumulation (16, 21), OpaR was shown to be required for CPS production and c-di-GMP accumulation in the *V. parahaemolyticus* strain BB22 (20, 22, 23). The absence of *opaR* in strain BB22 does not eliminate biofilm formation but rather affects the biofilm architecture, and potentially delays biofilm dispersal or detachment in the liquid-solid interface (22). Recent reports indicated that OpaR can repress biofilm formation and c-di-GMP accumulation in the *V. parahaemolyticus* strain RIMD2210663 (24). However, the involvement of OpaR in biofilm formation in strain RIMD2210633 was reported for one time point while for strain BB22 the kinetics of biofilm formation were evaluated.

Furthermore, the conditions of biofilm growth were different in both reports. For these reasons, it is unclear if the apparent discrepancies between these two strains are due to strain variability or the result of a bipolar role of OpaR in the regulation of biofilm formation, *cpsA* expression and c-di-GMP accumulation in response to environmental cues.

In a previous report we characterized a novel trigger phosphodiesterase in *V. parahaemolyticus* which we named TpdA (25). Trigger phosphodiesterases are c-di-GMP effectors that have enzymatic and regulatory activities that are typically anticorrelated (26). The enzymatic activity of TpdA anticorrelates with its ability to activate its own transcription (25). The ability of TpdA to promote its own transcription when c-di-GMP levels drop could accelerate c-di-GMP depletion or extend its duration. On the other hand, preventing TpdA expression might be necessary to set a c-di-GMP baseline that favors a rapid onset of biofilm formation in response to favorable conditions. The absence of TpdA has an effect on the intracellular levels of c-di-GMP but it has not been reported how this compares to the influence of other important PDEs such as ScrC and ScrG (27–29). These comparisons are key to understand how c-di-GMP can be controlled in a dynamic fashion.

Our main goal for this work was to further characterize the c-di-GMP-dependent regulation of *tpdA*. We hypothesized that OpaR, being a key modulator of c-di-GMP homeostasis, would regulate *tpdA* expression (23, 24). Our results uncovered a differential influence of OpaR-regulated PDEs (ScrC, ScrG and VP0117) on c-di-GMP accumulation and *tpdA* expression. We also observed a dominant c-di-GMP-degrading role for TpdA in planktonic cultures compared to ScrC, ScrG and VP0117. In contrast, ScrC played the central role as c-di-GMP degrader for the first 72 hours of growth over solid media until TpdA took over after 96 hours of growth. We also provide evidence that suggests that OpaR can act both as positive and negative modulator of c-di-GMP accumulation and *cpsA* expression under different growth conditions. This yin and yang nature of OpaR its likely enabled by its repertoire of regulatory targets that not only include PDEs but also DGCs and potentially dual function DGC-PDEs.

## Results

### The expression level of *tpdA* is altered by the absence of OpaR and OpaR-regulated phosphodiesterases

The activity of the P*_tpdA_* promoter is induced by a decrease in c-di-GMP accumulation, in a TpdA dependent manner (25). Although genetic evidence has suggested that TpdA is an activator of the P*_tpdA_* promoter, it is still unclear if this regulation is direct. We have speculated that other regulators might be involved in controlling the activity of the P*_tpdA_* promoter, but none has been characterized yet. Previous reports have shown that the master regulator of the quorum sensing response, OpaR, regulates several genes whose products are involved in the metabolism of c-di-GMP (24). A putative OpaR binding site was previously identified in the regulatory region of *tpdA* (23). To determine if the presence of OpaR affects the transcriptional activity of the P*_tpdA_* promoter we mobilized the transcriptional fusion *PtpdA-luxCDABE*, assembled into the pBBRlux plasmid, to the wild-type (WT) strain and to a *ΔopaR* mutant strain. In this assay, light production is proportional to the activity of the P*_tpdA_* promoter. Our results revealed a 55%, 79%, 166% and 126% increase in activity of the P*_tpdA_* promoter in the *ΔopaR* mutant strain compared to the WT strain after 4, 6, 8 and 24 hours of growth, respectively (Fig. 1C-F). These results strongly suggest that OpaR negatively regulates the expression of *tpdA*.

**Fig. 1.**
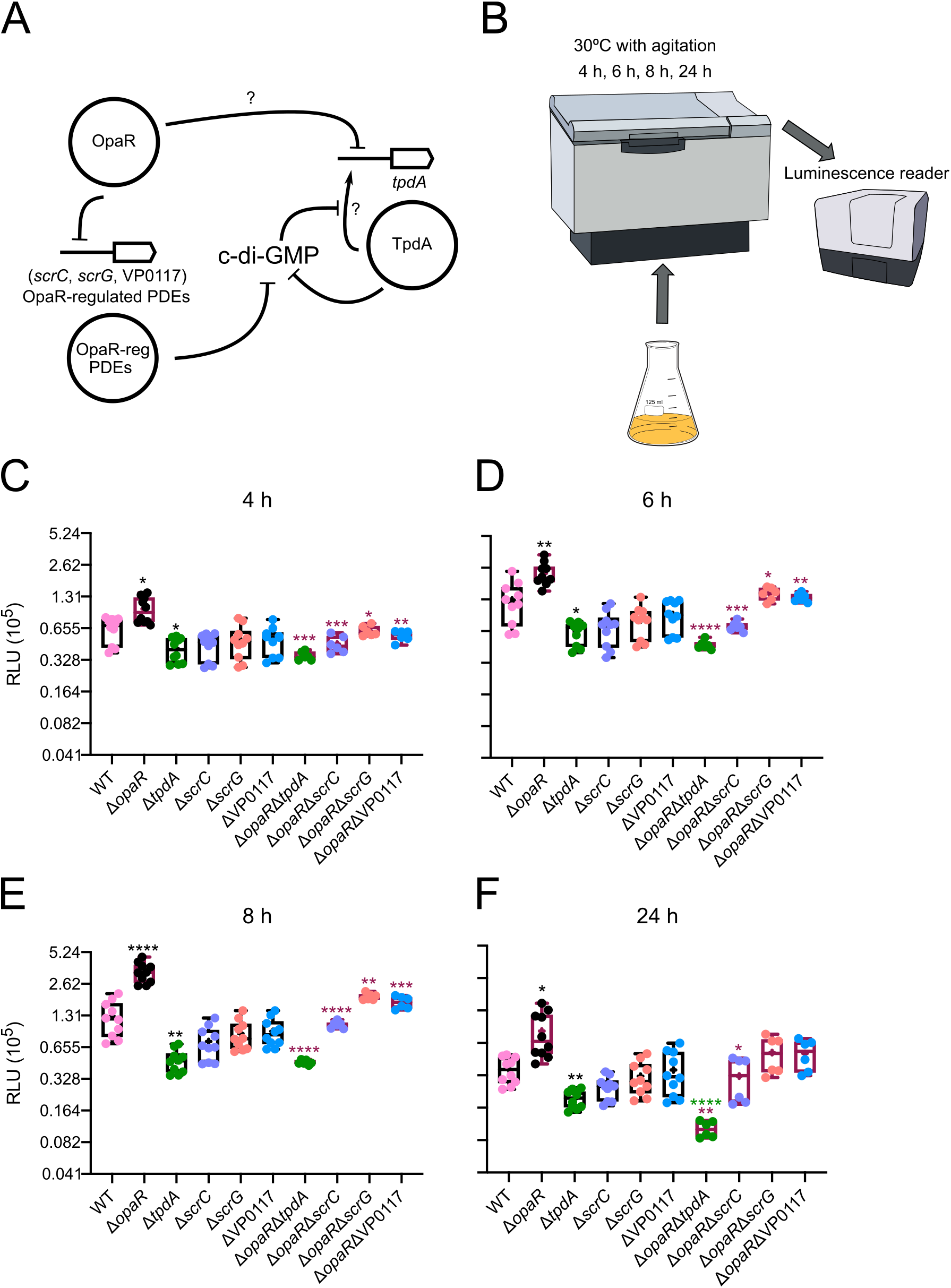
The expression of *tpdA* is oppositely influenced by OpaR and OpaR-regulated PDEs. A) Schematic representation of a proposed model for c-di-GMP modulation through OpaR and its regulated PDEs, and the potential effect of this modulation on *tpdA* expression. Arrows represent positive transcriptional regulation, while T connectors represent negative transcriptional regulation or enzymatic degradation of c-di-GMP. Question marks indicate that it is unknown whether the regulation is direct or indirect. B) Schematic representation of the experimental procedure. Cells were grown as planktonic cultures with agitation at 30 °C. A minimum of six independent biological samples were analyzed, and experiments were done at least twice. C,D,E,F) Box plots representing the expression data, in Relative Light Units (RLU), of the transcriptional fusion *PtpdA-luxCDABE* in different genetic backgrounds at 4 time points. Means were compared using a Brown-Forsythe and Welch ANOVA test followed by a Dunett’s T3 multiple comparison test to compare directly to the mean of either the WT strain or the mutant strains *ΔopaR* or *ΔtpdA*. Black, maroon and green asterisks indicate a statistical difference compared to WT, *ΔopaR* or *ΔtpdA*, respectively. Adjusted P value * ≤ 0.05, ** ≤ 0.01 *** ≤ 0.001 **** ≤ 0.0001.

We have previously reported that the activity of P*_tpdA_* is induced when an active PDE is overproduced (25). OpaR regulates the production of multiple PDEs, with ScrC and ScrG among the most well characterized (Fig. 1A) (27–29). The expression of *scrC* is regulated negatively by OpaR, while the expression of *scrG* appears to be positively influenced by OpaR (23, 24). We decided to analyze if ScrC, ScrG and also VP0117, another putative c-di-GMP metabolizing enzyme negatively regulated by OpaR (24), influence the activity of the P*_tpdA_* promoter. The absence of *tpdA* resulted in a 35%, 50%, 62% and 47% decrease in the activity of the P*_tpdA_* promoter compared to the WT strain after 4, 6, 8 and 24 hours of growth, respectively (Fig. 1CDEF). The absence of *scrC, scrG and* VP0117 did not significantly affect the activity of the P*_tpdA_* promoter (Fig. 1C-F). Next, we investigated the contribution of the OpaR-regulated PDEs to the induction of the P*_tpdA_* promoter observed in the *ΔopaR* genetic background. Our results revealed that the absence of *opaR* cannot induce the activity of the P*_tpdA_* promoter when *tpdA* is also absent *(ΔopaR ΔtpdA)* (Fig. 1C-F). We also observed that after 24 hours of growth the activity of the P*_tpdA_* promoter is lower in the double *ΔopaR ΔtpdA* mutant strain than in the single *ΔtpdA* mutant strain (Fig. 1F). This would suggest that the activity of this promoter is also positively influenced by the presence of OpaR under certain conditions. The activity of the P*_tpdA_* promoter in the *ΔopaR ΔscrC* mutant strain decreased 54%, 69%, 69%, and 62% after 4, 6, 8, and 24 hours of growth, respectively, compared to the *ΔopaR* mutant strain (Fig. 1C-F). Since OpaR negatively regulates *scrC*, our results would suggest that ScrC is a key contributor to the upregulation of *tpdA* in the *ΔopaR* genetic background, although not the only one. The contribution of ScrG and VP0117 to the elevation of P*_tpdA_* activity in the *ΔopaR* mutant strain was modest (Fig. 1C-F). After 8 hours of growth the double mutants the *ΔopaR ΔscrG* and *ΔopaR* ΔVP0117 showed a 42% and 49% decrease in P*_tpdA_* activity, respectively, compared to the *ΔopaR* mutant strain (Fig. 1E). Together, our results revealed a hierarchical control of the activity of the P*_tpdA_* promoter by OpaR-regulated PDEs (TpdA>ScrC>VP0117>ScrG).

### OpaR has opposite effects on c-di-GMP accumulation at early and late stages of planktonic growth

As mentioned above, the activity of the P*_tpdA_* promoter can be induced by a decrease in c-di-GMP levels (25). Based on this antecedent we speculated that the induction of the activity of the P*_tpdA_* promoter in the *ΔopaR* genetic background could be due to a decrease in c-di-GMP levels. Furthermore, we also hypothesized that the hierarchical regulation of the P*_tpdA_* promoter by ScrC, VP0117 and ScrG could be explained by differences in the levels of c-di-GMP in genetic backgrounds lacking these PDEs. To evaluate c-di-GMP levels in the genetic backgrounds of interest we used a genetic biosensor previously shown to be able to detect differences in c-di-GMP levels in *Vibrio cholerae, Vibrio parahaemolyticus* and *Vibrio fischeri* (25, 30, 31). In this c-di-GMP biosensor the production of two fluorescent proteins, Amcyan and TurboRFP is controlled by a constitutive promoter (30). While production of Amcyan is not directly affected by c-di-GMP levels, the production of TurboRFP is directly controlled by two c-di-GMP riboswitches in tandem (30). Binding of c-di-GMP to the riboswitches favors the production of TurboRFP. Hence, the fluorescence of TurboRFP is proportional to the level of c-di-GMP. The fluorescence of Amcyan is used to normalize for differences in the abundance of the biosensor or the activity of its promoter. Our results revealed that the level of c-di-GMP in the absence of *opaR (ΔopaR)* decreases 27%, 43%, and 48% after 4, 6 and 8 hours of planktonic growth, respectively, compared to the levels in the WT strain (Fig. 2B-D). In contrast, after 24 hours of growth the c-di-GMP levels in the *ΔopaR* mutant strain and the WT strain were not significantly different (Fig. 2E). These results could be interpreted as OpaR being responsible for maintaining the baseline levels of c-di-GMP during the first 8 hours of growth, and either stops contributing or inhibits its accumulation at some point between the 8 and 24 hours of growth. The dual nature of OpaR in terms of its role in controlling c-di-GMP homeostasis becomes evident when the absence of this regulator is combined with the absence of the PDE TpdA. First, the *ΔtpdA* mutant strain, as previously reported, showed approximately a 20% increase in c-di-GMP accumulation compared to the WT strain at all time points evaluated. This was the only PDE that contributed to c-di-GMP homeostasis under our experimental conditions either in the WT or *ΔopaR* genetic background (Fig. 2). The *ΔopaR ΔtpdA* double mutant showed an intermediate phenotype in terms of c-di-GMP accumulation compared to the single mutants *ΔopaR* and *ΔtpdA*, at each time point analyzed up to the 8 hours of growth. Interestingly, at the 24-hour time point the c-di-GMP levels in the *ΔopaR ΔtpdA* mutant strain showed a 74% increase compared to the levels in the WT strain, which was higher than the increase in c-di-GMP accumulation in the single mutant *ΔtpdA*. The additive effect of the combined absence of *opaR* and *tpdA* on c-di-GMP accumulation compared to the WT strain and the single mutants strongly suggests that OpaR prevents c-di-GMP accumulation during the late stationary phase of growth, at least when TpdA is absent.

**Fig. 2.**
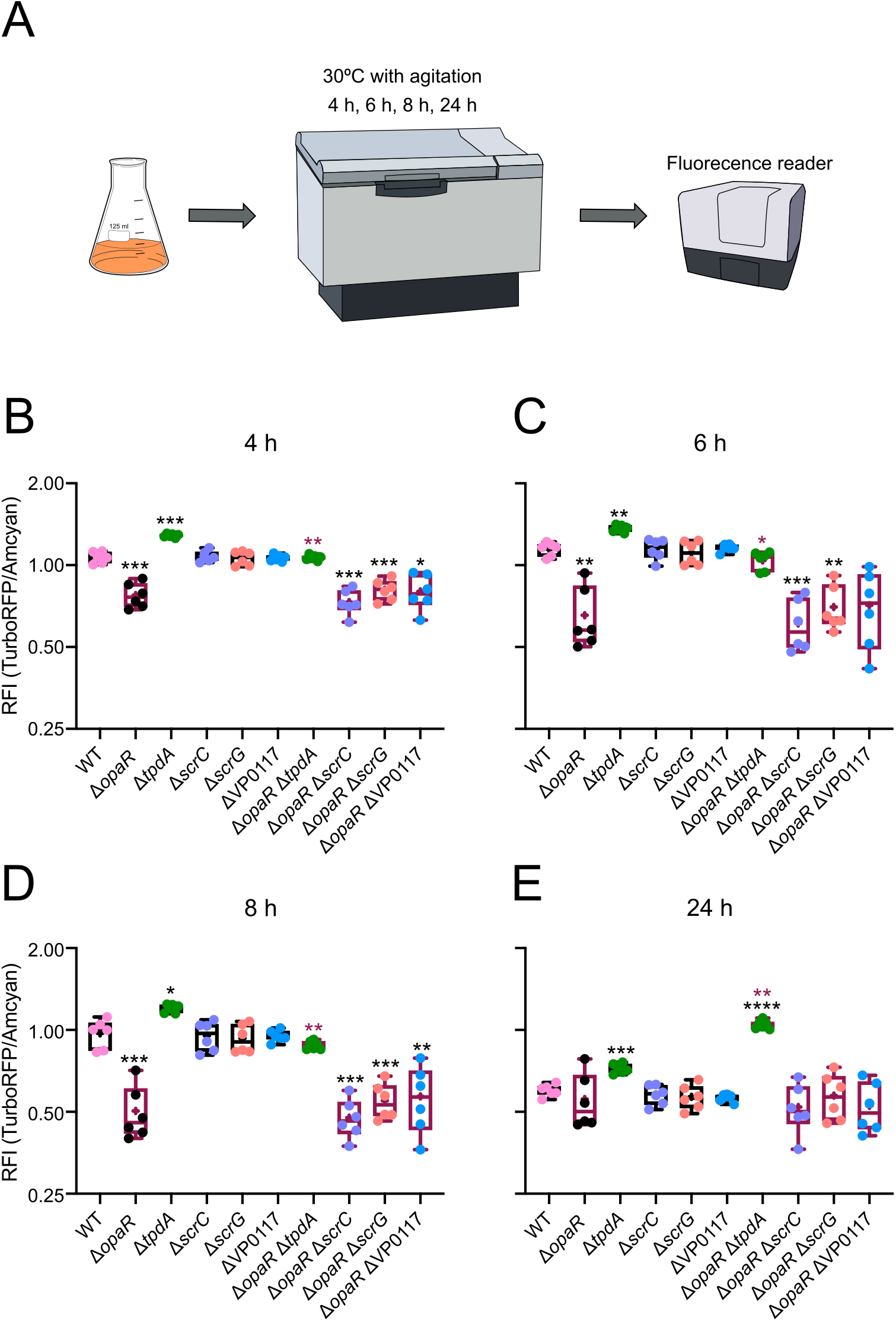
OpaR and TpdA are important modulators of c-di-GMP homeostasis in planktonic cultures. A) Schematic representation of the experimental procedure. Cultures were grown planktonically with agitation at 30 °C. At least six independent biological samples were analyzed across three independent experiments. B,C,D,E) Box plots representing the values of Relative Fluorescence Intensity (RFI). The RFI values are proportional to the level of c-di-GMP in the cultures at 4 different time points. Means were compared using a Brown-Forsythe and Welch ANOVA test followed by a Dunett’s T3 multiple comparison test to compare directly to the mean of either the WT strain or the *ΔopaR* mutant strain. Black and maroon asterisks indicate a statistical difference compared to WT or *ΔopaR*, respectively. Adjusted P value * ≤ 0.05, ** ≤ 0.01 *** ≤ 0.001 **** ≤ 0.0001.

Our observations suggest that TpdA is one of the most important contributors to the maintenance of c-di-GMP levels in planktonic cultures. The contribution of ScrC, ScrG and VP0117 might be cumulative but is also possible that other OpaR-regulated PDEs and DGCs contribute to the maintenance of c-di-GMP levels.

### OpaR negatively regulates the expression of *tpdA* by altering the global c-di-GMP pool

Since we had demonstrated that the absence of *opaR* results in a decrease in c-di-GMP accumulation compared to the WT strain (Fig. 2), we next asked if the change in c-di-GMP levels in this genetic background could be the main cause for the induction of the activity of the P*_tpdA_* promoter. We artificially elevated the c-di-GMP levels in the *ΔopaR* mutant strain by introducing a plasmid that enables the overexpression of *cdgF* (VCA0956), whose product is an active DGC from *V. cholerae* (12). To determine the magnitude of c-di-GMP increase resulting from the overexpression of *cdgF*, we introduced a compatible plasmid that has the c-di-GMP biosensor described above (pDZ119) (25). As a control, we introduced the transcriptional fusion or the c-di-GMP biosensor together with the empty expression plasmid into the WT strain and the *ΔopaR* mutant strain to have conditions were *cdgF* is not overexpressed. The overproduction of CdgF in the *ΔopaR* mutant strain resulted in c-di-GMP accumulation levels similar to those observed in the WT strain (Fig. 3A). This increase in c-di-GMP levels was enough to eliminate the induction of the P*_tpdA_* promoter (Fig. 3B). This result strongly suggests that the negative effect of OpaR over the activity of the P*_tpdA_* promoter is mainly due to the ability of this regulator to dictate the baseline level of c-di-GMP in planktonic cultures.

**Fig. 3.**
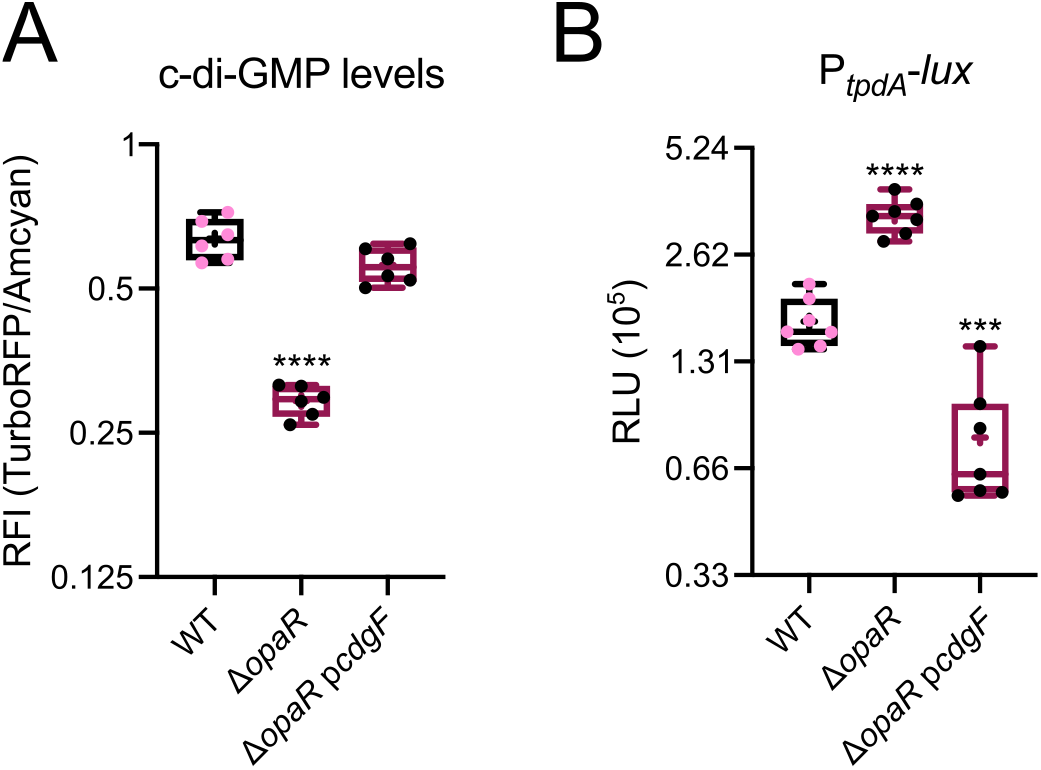
An increase in c-di-GMP levels eliminates the upregulation of *tpdA* observed in the absence of *opaR*. A) Box plots representing the c-di-GMP levels measured with the c-di-GMP genetic reporter in cells grown for 8 hours in agitation at 30 °C in the presence of 0.1 mM IPTG. C-di-GMP levels are expressed as RFI values. B) Box plots represent the expression data, in RLU, of the transcriptional fusion *PtpdA-luxCDABE* in different genetic backgrounds after 8 hours of growth in agitation at 30 °C in the presence of 0.1 mM IPTG. Three independent biological samples were analyzed in two separate experiments. Means were compared using a One-way ANOVA test followed by a Dunett’s multiple comparison test to compare directly to the mean of the WT strain. Asterisks indicate a statistical difference compared to WT. Adjusted P value *** ≤ 0.001 **** ≤ 0.0001.

### The expression of *tpdA* in cells growing over solid media showed an altered dynamic sensitivity to the absence of OpaR or ScrC compared to cells grown in liquid cultures

Our results so far indicate that OpaR has a negative effect on *tpdA* expression, and that the regulatory mechanism involves the manipulation of c-di-GMP levels in part through the control of ScrC abundance. ScrC production is directly regulated by OpaR but its activity as a PDE is also influenced by the abundance of an autoinducer-like molecule produced by ScrA (28). It has been speculated that the accumulation of this autoinducer-like signal is favored by a surface-adhered lifestyle (28). These antecedents led us to evaluate the influence of OpaR, ScrC and TpdA on the transcriptional dynamics of the P*_tpdA_* promoter in cells growing on solid media (LB-agar). We analyzed the activity of the transcriptional fusion *PtpdA-luxCDABE* in the WT, *ΔopaR, ΔtpdA, ΔscrC, ΔopaR ΔtpdA*, and *ΔopaR ΔscrC* genetic backgrounds in cells growing over LB-agar for 24, 48, 72 and 96 hours at 30 °C (Fig. 4A).

**Fig. 4.**
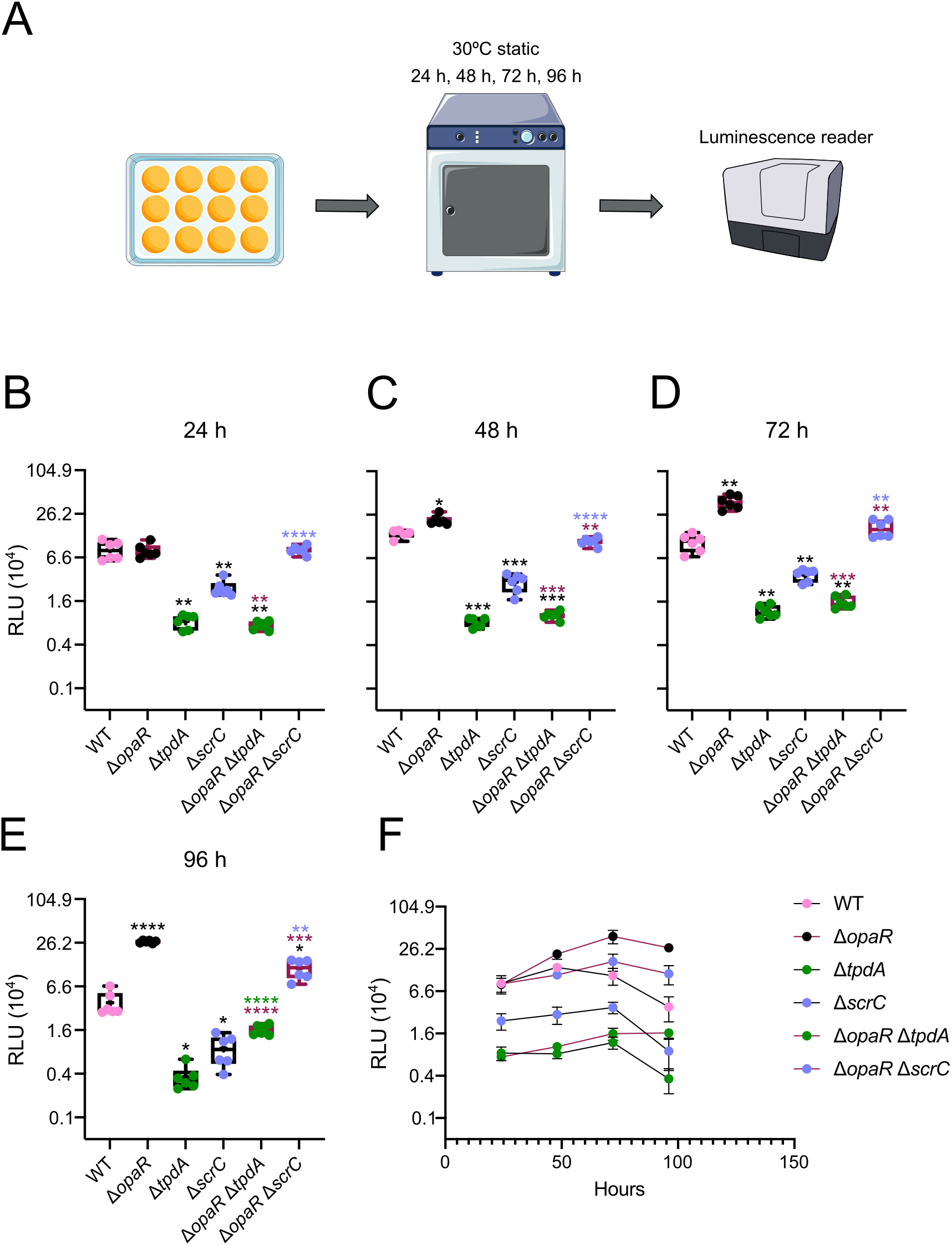
The influence of OpaR over the expression of *tpdA* increases over time in cells grown on solid media. A) Schematic representation of the experimental procedure. Cells were grown statically over LB agar with 15 μg/ml Chl at 30 °C. At least five independent biological samples were analyzed across two independent experiments. B,C,D,E) Box plots representing the expression data, in RLU, of the transcriptional fusion *PtpdA-luxCDABE* in different genetic backgrounds at 4 time points. F) XY graph of the data plotted in B, C, D, and E showing the mean values of RLU for each strain at each time point tested. Error bars represent the standard deviation. Means were compared using a Brown-Forsythe and Welch ANOVA test followed by a Dunett’s T3 multiple comparison test to compare directly to the mean of either the WT strain, the *ΔopaR, ΔtpdA*, or Δ*scrC* mutant strains. Black, maroon, green, or orchid asterisks indicate a statistical difference compared to WT, *ΔopaR, ΔtpdA*, or Δ*scrC*, respectively. Adjusted P value * ≤ 0.05, ** ≤ 0.01 *** ≤ 0.001 **** ≤ 0.0001.

The activity of the P*_tpdA_* promoter in the WT strain increased 70% between the 24-hour and 48-hour time points (Fig. 4B-C, F). After this peak in expression, the activity of the P*_tpdA_* promoter started to decline and ended up showing a 54% decrease compared to the initial measurement in the growth curve (24 h vs 96 h) (Fig. 4B-F). In contrast to what we observed in planktonic cultures, the absence of *opaR* did not affect the activity of the P*_tpdA_* promoter after 24 hours of growth (Fig. 4B). However, we observed a 55%, 264% and 593% increase in the activity of the P*_tpdA_* promoter in the *ΔopaR* mutant strain compared to the WT strain after 48, 72 and 96 hours of growth, respectively (Fig. 4C-E). The peak of activity of the P*_tpdA_* promoter in the *ΔopaR* mutant strain was reached after 72 hours of growth (Fig. 4F). This could suggest that OpaR is responsible for limiting the expression of *tpdA* at later time points during the growth over LB-agar.

The activity of the P*_tpdA_* promoter decreased 90% and 71% in the *ΔtpdA* and *ΔscrC* mutant strains, respectively, after 24 hours of growth, compared to the WT strain (Fig. 4B), and remained lower than in the WT for the next time points. In planktonic cultures we did not observe an effect on *tpdA* expression in the *ΔscrC* single mutant strain at any time point, hence it appears that ScrC plays a more determinant role in controlling the expression of *tpdA* in cells grown on solid media.

In planktonic cultures we observed that the absence of *tpdA* was dominant over the absence of *opaR* with regards to their effect on the activity of the P*_tpdA_* promoter (Fig. 1). Our data also suggested that at later stages of the growth curve, both genes could play a positive role on the activity of the P*_tpdA_* promoter (Fig. 1F). In cells grown over LB-agar, we also observed a dominant effect of the absence of *tpdA* over the absence of *opaR*. However, in contrast to what we observed in shaking cultures, the activity of the P*_tpdA_* promoter in the *ΔopaR ΔtpdA* mutant strain showed a 350% increase compared to the *ΔtpdA* mutant strain in the last time point measured (96 hours). This could suggest that under these conditions of growth, the activity of the P*_tpdA_* promoter can be induced independently of TpdA in the absence of OpaR.

The activity of the P*_tpdA_* promoter in the *ΔopaR ΔscrC* strain was not statistically different compared to the WT or the *ΔopaR* strains up to 24 hrs. From 48 hrs to 72 hrs of growth the activity levels of P*_tpdA_* in the *ΔopaR ΔscrC* strain remain similar to the WT strain but decrease in comparison to the *ΔopaR* strain, and at the 96 hrs time point they are lower when compared to both the WT and the *ΔopaR* strain. The intermediate effect on *tpdA* expression of the combined versus single absence of *opaR* and *scrC* genes from the 48 hrs to the 96 hrs time points supports the notion that multiple OpaR-regulated genes are involved in controlling the production of the trigger phosphodiesterase TpdA.

### ScrC and TpdA exchange the dominant role on c-di-GMP degradation after the transition from the early to late stage of growth over LB-agar, in the absence of OpaR

Our observations regarding the activity of the P*_tpdA_* promoter in cells grown over LB-agar prompted us to analyze the dynamics of c-di-GMP accumulation under the same conditions of growth (Fig. 5A). C-di-GMP levels increased 197% between 24 hours and 72 hours of growth in the WT strain and remained relatively stable afterwards (Fig. 5). The absence of *opaR* resulted in a 73%, 90%, 92%, 89% and 83% decrease in c-di-GMP levels after 24, 48, 72, 96 and 120 hours of growth, respectively, compared to the WT strain. This result strongly suggests that OpaR plays a key role in promoting c-di-GMP accumulation in cells growing over LB-agar. We previously showed that the expression of *tpdA* is not altered in the *ΔopaR* mutant strain compared to the WT strain after 24 hours of growth over LB-agar (Fig. 4B), and yet we are reporting a 73% decrease in c-di-GMP levels in the *ΔopaR* mutant strain compared to the WT strain at the 24-hour time point. It is possible that factors other than the level of c-di-GMP influence the activity of the P*_tpdA_* promoter in cells grown on LB-agar during the initial hours of growth.

**Fig. 5.**
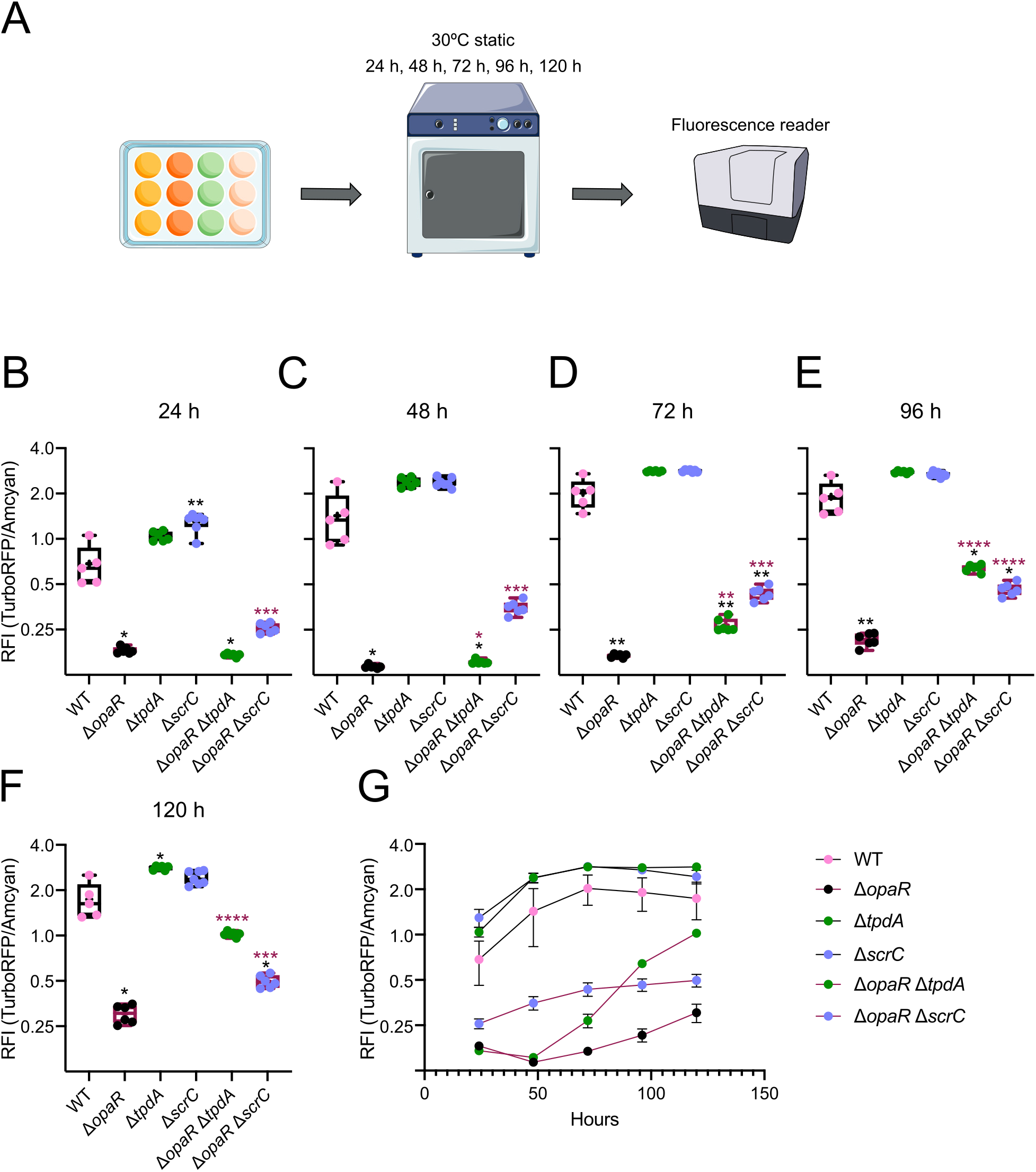
The OpaR regulated PDEs TpdA and ScrC act at different time windows to control c-di-GMP metabolism in cells growing on solid media. A) Schematic representation of the experimental procedure. Cells were grown statically over LB agar with 15 μg/ml Gen at 30 °C. At least 5 independent biological samples were analyzed across three independent experiments. B,C,D,E,F) Box plots representing the values of RFI, which are proportional to the level of c-di-GMP in the colonies at 5 different time points. G) XY graph of the data plotted in B, C, D, E and F showing the mean values of RFI for each strain at each time point tested. Error bars represent the standard deviation. Means were compared using a Brown-Forsythe and Welch ANOVA test followed by a Dunett’s T3 multiple comparison test to compare directly to the mean of either the WT strain or the *ΔopaR* mutant strain. Black and maroon asterisks indicate a statistical difference compared to WT or *ΔopaR*, respectively. Adjusted P value * ≤ 0.05, ** ≤ 0.01 *** ≤ 0.001 **** ≤ 0.0001.

Although the absence of *opaR* had a profound effect on c-di-GMP accumulation compared to the WT strain, we observed a 66% increase in c-di-GMP levels in the *ΔopaR* mutant strain when comparing the 24-hour and 120-hour time points. Since the trend in c-di-GMP accumulation is the opposite in the WT and the *ΔopaR* mutant strain after 72 hours of growth, we speculate that the role of OpaR as a regulator of c-di-GMP may switch from positive to negative at later growth stages in cells adhered to LB-agar. It is also possible that an OpaR-independent control mechanism is responsible of promoting c-di-GMP accumulation after a prolonged period where the levels of this second messenger are below the “expected” baseline.

The differences in c-di-GMP accumulation between the WT strain and the single mutants in *scrC* and *tpdA* were more evident at the 24-hour and 120-hour time points, respectively (Fig. 5). After 24 hours of growth, we observed a 90% increase in c-di-GMP accumulation in the Δ*scrC* mutant strain compared to the WT strain, while after 120 hours of growth we observed a 62% increase in c-di-GMP levels in the *ΔtpdA* mutant strain compared to the WT strain. The involvement of TpdA and ScrC in controlling c-di-GMP homeostasis in cells growing over LB-agar was more evident when comparing c-di-GMP levels between the *ΔopaR* single mutant and the double mutants *ΔopaR ΔtpdA* and *ΔopaR ΔscrC*. The c-di-GMP levels in the *ΔopaR* and *ΔopaR ΔtpdA* mutant strains were very similar after 24 and 48 hours of growth. After 72, 96 and 120 hours of growth we observed a 60%, 199% and 238% increase in c-di-GMP accumulation, respectively, in the *ΔopaR ΔtpdA* mutant strain compared to the *ΔopaR* mutant strain. This result suggests that TpdA does not play a dominant role in depleting c-di-GMP levels in the absence of OpaR in the early time points (24 and 48 hours), but its presence is very influential in controlling c-di-GMP accumulation at later time points. On the other hand, we observed an increase in c-di-GMP levels in the *ΔopaR ΔscrC* mutant strain compared to the *ΔopaR* mutant strain at all the time points analyzed. This could indicate that ScrC plays a dominant role in controlling c-di-GMP levels in the absence of OpaR. However, at 96 and 120 hours of growth the increase in c-di-GMP levels was higher in the *ΔopaR ΔtpdA* mutant strain compared to the *ΔopaR ΔscrC* mutant strain, suggesting that the dominant influence in controlling c-di-GMP levels in cells growing over LB-agar in the absence of OpaR switches from ScrC to TpdA at the 96-hour time point (Fig. 5D-F).

In summary, our data suggest that the OpaR-regulated PDEs ScrC and TpdA act at different stages of colony-growth to control c-di-GMP levels. The differences in ScrC and TpdA influence over c-di-GMP homeostasis are more evident in the absence of OpaR perhaps due to the negative effect of this regulator on the expression of both *scrC* and *tpdA*. It is also clear from these results that OpaR can control c-di-GMP accumulation in cells grown on LB-agar through additional regulators besides ScrC and TpdA.

### OpaR, ScrC and TpdA affect the dynamics of *cpsA* expression in cells growing over LB-agar

Since c-di-GMP plays a key role in controlling the expression of *cpsA* (25, 29, 32, 33), we next analyzed if the drop or elevation in c-di-GMP levels experienced by our strains of interest when grown over LB-agar, results in altered expression of *cpsA*. For the analysis of *cpsA* expression we used a previously reported transcriptional fusion with the P_*cpsA*_ promoter fused to the reporter genes *luxCDABE* (25) (Fig. 6A). Our results revealed that the expression of *cpsA* is negatively affected by the absence of OpaR after 24, 48, 72 hours of growth (Fig. 6B-D) and positively impacted by the absence of ScrC after 48 and 72 hours of growth (Fig. 6C-D). The expression of *cpsA* in the *ΔopaR ΔtpdA* mutant strain showed similar levels to those observed in the *ΔopaR* mutant strain from the 24-to the 72-hour time point (Fig. 6B-D). After 96 hours of growth the expression of *cpsA* in the *ΔopaR ΔtpdA* mutant strain showed a 143% and 165% increase in expression when compared to the *ΔopaR* and *ΔtpdA* mutant strains, respectively (Fig. 6E). This suggest that after 96 hours of growth the combined absence of *opaR* and *tdpA* has a stronger positive effect on *cpsA* expression than when these genes are missing individually. The expression of *cpsA* in the *ΔopaR ΔscrC* double mutant strain showed a peculiar behavior when compared to any of the other strains analyzed (Fig. 6F). After 24 hours of growth, the level of activity of the P_*cpsA*_ promoter in the *ΔopaR ΔscrC* mutant strain was closer to the level observed in the WT strain (Fig. 6B). Then, after 48 hours of growth, the activity of the P_*cpsA*_ promoter in the *ΔopaR ΔscrC* strain dropped to levels reached in the *ΔopaR* mutant strain only to spike to levels of activity observed in the *ΔscrC* mutant strain after 72 hours of growth (Fig. 6C-D). By looking at the 24- and 72-hour time points, we would conclude that ScrC is either partially (at 24 hours) or completely dominant over OpaR with regards to *cpsA* regulation. However, at the 48-hour time point the positive role of OpaR over *cpsA* expression is dominant over the negative regulation exerted by ScrC. These results suggest that the requirement for the presence of OpaR to activate *cpsA* expression in the absence of ScrC is variable and depends on unknown regulatory factors.

**Fig. 6.**
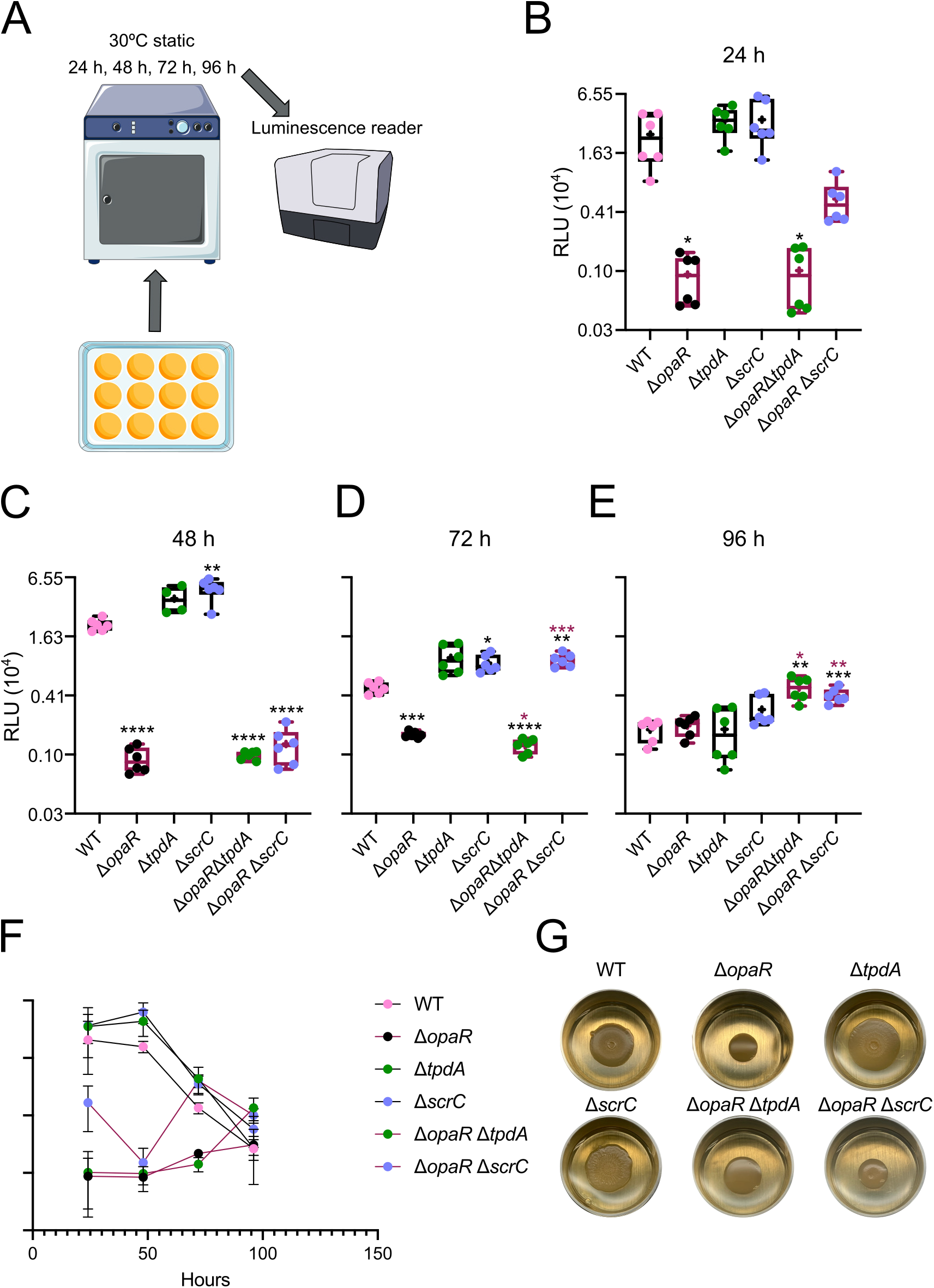
OpaR and its regulated PDEs contribute to control the expression of *cpsA* over time in cells grown on solid media. A) Schematic representation of the experimental procedure. Cells were grown statically over LB agar with 15 μg/ml Chl at 30 °C. At least 4 independent biological samples were analyzed across two independent experiments. B,C,D,E) Box plots representing the expression data, in RLU, of the transcriptional fusion *PcpsA-luxCDABE* in different genetic backgrounds at 4 time points. F) XY graph of the data plotted in B, C, D, and E showing the mean values of RLU for each strain at each time point tested. Error bars represent the standard deviation. Means were compared using a Brown-Forsythe and Welch ANOVA test followed by a Dunett’s T3 multiple comparison test to compare directly to the mean of either the WT strain, or the *ΔopaR* mutant strain. Black and maroon asterisks indicate a statistical difference compared to WT and *ΔopaR*, respectively. Adjusted P value * ≤ 0.05, ** ≤ 0.01 *** ≤ 0.001 **** ≤ 0.0001. G) Pictures of representative spot-colonies used in the assay. These colonies were grown for 48 h at 30°C in 12-well plates filled with solid media.

We also documented representative images of the spot-colony morphology of the strains of interests carrying the *PcpsA-luxCDABE* transcriptional fusion plasmid after 48 hours of growth (Fig. 6G). The morphology of the *ΔopaR* and the *ΔscrC* mutant strains was clearly different from the morphology of the WT strain (Fig. 6G). The spot-colonies of the *ΔopaR* mutant strain were more constrained and smoother compared to the WT strain (Fig. 6G). On the other hand, the spot-colonies of the *ΔscrC* mutant strain were more spread and rugose compared to the WT strain (Fig. 6G). The spot-colonies of the *ΔtpdA* mutant strain were also more spread compared to the WT strain but were not as rugose as those of the *ΔscrC* mutant strain (Fig. 6G). The morphology of the spot-colonies made by the double mutants *ΔopaR ΔtpdA* and *ΔopaR ΔscrC* resemble that of the *ΔopaR* mutant strain (Fig. 6G). The spot colonies of the *ΔopaR ΔtpdA* mutant strain were more spread compared to the *ΔopaR* mutant strain and similarly smooth (Fig. 6G). On the other hand, the spot colonies of the *ΔopaR ΔscrC* mutant strain were of similar size compared to the *ΔopaR* mutant strain but less smooth (Fig. 6G).

Our data suggest that the control exerted by OpaR over c-di-GMP accumulation in cells growing attached to solid media influences *cpsA* expression and that such control is dynamic, switching from positive to negative after 48 hours of growth.

### OpaR is capable of inhibiting biofilm formation and *cpsA* expression

Our results so far support the role of OpaR as a positive regulator of *cpsA* expression and in consequence of biofilm matrix production in cells growing over solid media (19, 20, 22, 23, 34). However, when growing overnight cultures of the *ΔopaR* mutant strain under shaking conditions we noticed the formation of a thick biofilm on the surface of glass test tubes. Thus, we decided to make a quantitative analysis of the level of *cpsA* expression and c-di-GMP accumulation in this type of biofilm. For these experiments we used strains harboring the plasmid pDZ46 or pFY4535, which have the *PcpsA-luxCDABE* transcriptional fusion or the c-di-GMP biosensor, respectively. Our results revealed a 210% and 300% increase in biofilm formation under shaking conditions over a glass surface in the *ΔopaR* mutant strain harboring pDZ46 or pFY4535, respectively, compared to the WT strain (Fig. 7A-D). The *ΔtpdA* mutant strain made a similar amount of biofilm in comparison to the WT strain, while the *ΔscrC* mutant strain showed an increase in biofilm formation compared to the WT strain but only when comparing strains harboring the pDZ46 plasmid (Fig. 7A-D).

**Fig. 7.**
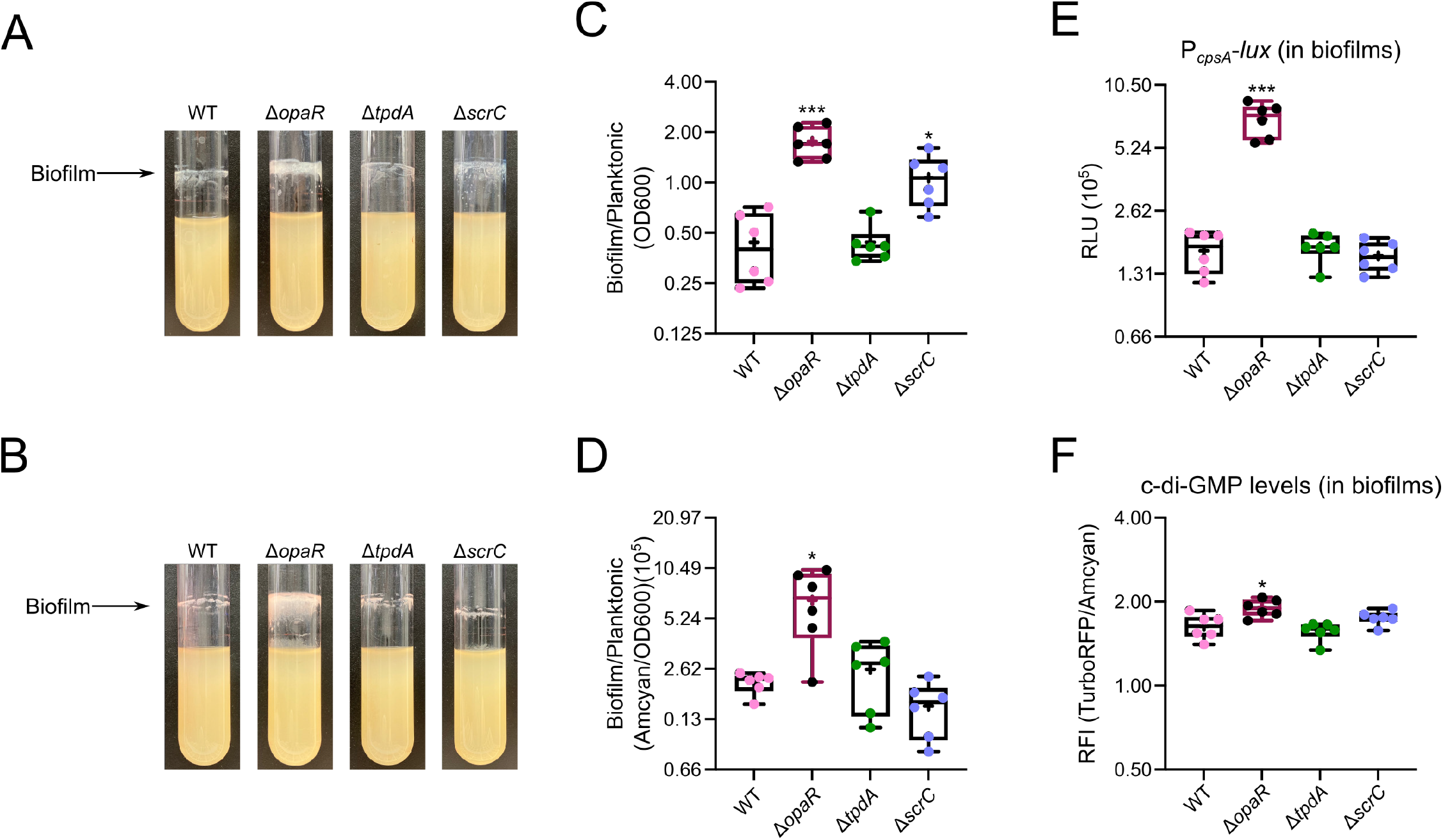
OpaR limits biofilm formation and *cpsA* expression in biofilms grown over glass in agitation. A,B) Representative pictures of 24-hour biofilms grown in shaking conditions over the glass surface of test tubes at 30 °C, from strains harboring the pBBRlux-P_*cpsA*_ plasmid (A) or the pFY4535 plasmid (B). C,D) Box plots representing the quantification of biofilm formation from strains harboring the pBBRlux-P_*cpsA*_plasmid (C) or the pFY4535 plasmid (D). E) Box plots representing the expression data, in RLU, of the transcriptional fusion *PcpsA-luxCDABE* in different genetic backgrounds. Biofilms expressing *PcpsA-luxCDABE* were harvested after 24 hours of growth in agitation at 30 °C. F). Box plots representing the c-di-GMP levels, expressed as RFI values, measured with the c-di-GMP genetic reporter in biofilms grown in agitation at 30 °C for 24 hours. Means were compared using a Brown-Forsythe and Welch ANOVA test followed by a Dunett’s T3 multiple comparison test to compare directly to the mean of the WT strain. Asterisks indicate a statistical difference compared to WT. Adjusted P value * ≤ 0.05, *** ≤ 0.001.

In terms of *cpsA* expression in biofilm cells, we found that the *ΔopaR* mutant strain had a 326% increase compared to the WT strain (Fig. 7E). On the other hand, the level of expression of *cpsA* in the *ΔtpdA* and *ΔscrC* mutant strains was not different to the level observed in the WT strain (Fig. 7E). This would suggest that under this growth conditions and time point analyzed, the expression of *cpsA* is strongly inhibited by OpaR but not by ScrC or TpdA. We also found a 16% increase in c-di-GMP accumulation in biofilms formed by the *ΔopaR* mutant strain compared to the WT strain. (Fig. 7F). It is unclear if such a minor elevation in c-di-GMP levels can explain the more profound difference in the amount of biofilm formation and the levels of expression of *cpsA* between the WT and *ΔopaR* mutant strain. It is possible that the change in c-di-GMP levels in the *ΔopaR* mutant strain under this condition is transient, while the resulting change in *cpsA* expression is more stable. As will be discussed later, OpaR is likely a direct regulator of *cpsA* expression and other biofilm related genes (23, 35, 36). Hence, we could also speculate that under certain circumstances OpaR could negatively regulate biofilm formation even in the absence of altered c-di-GMP levels. We next evaluated if under our experimental conditions the *ΔopaR* mutant strain had an altered biofilm phenotype compared to the WT strain under static growth over a plastic surface (PVC), as previously reported in the genetic background of strain BB22 (22). We analyzed the kinetics of biofilm formation under static conditions in PVC wells of the WT strain, the *ΔopaR, ΔtpdA, ΔscrC, ΔopaR ΔtpdA*, and *ΔopaR ΔscrC* mutant strains, and control strains with low *(ΔcpsR)* or high biofilm formation capacity *(ΔcpsS)* (Fig. 8) (20). The WT strain showed a peak of biofilm formation after 4 hours of growth in LB at 30 °C; after this time point the biofilm started to disperse and/or detach (Fig. 8). The *ΔopaR* mutant strain showed similar levels of biofilms formation compared to the WT strain after 4 hours of growth. However, at the 6-hour time point the *ΔopaR* mutant strain showed a 517% increase in biofilm formation compared to the WT strain. The 6 hour-time point marked the peak of biofilm formation for the *ΔopaR* mutant strain. While in the WT strain biofilm formation decreased 62% and 86% after 6 and 8 hours of growth compared to the 4-hour time point, in the *ΔopaR* mutant strain biofilm formation remained stable from the 6- to the 8-hour time point. It can be proposed that the absence of *opaR* partially prevents biofilm dispersal, perhaps by extending the time window of expression of biofilm related genes. These results are similar to the previously reported observations made in the *V. parahaemolyticus* strain BB22 (22). The individual absence of *tpdA* or *scrC* does not affect biofilm formation compared to the WT strain. When, *tpdA* was eliminated together with *opaR* the biofilm formation phenotype resembled that of the *ΔopaR* mutant strain. Similarly, when *scrC* was eliminated together with *opaR*, we observed only a minor increase in biofilm formation compared to the *ΔopaR* mutant strain. These results suggest that TpdA and ScrC do not play a major role in controlling biofilm formation under these conditions.

**Fig. 8.**
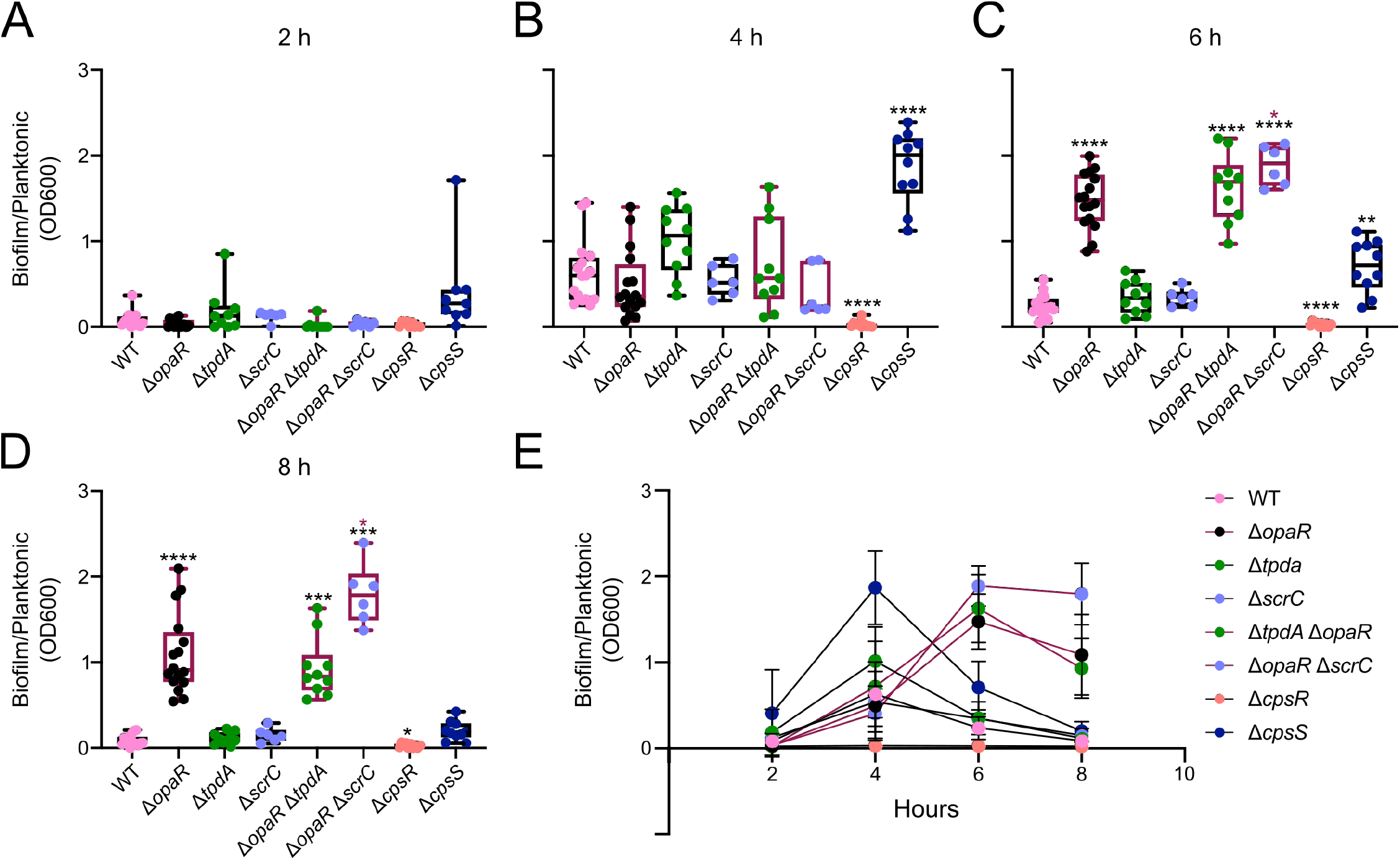
OpaR plays a key role in controlling the kinetic of biofilm formation over a PVC surface. A, B, C, D) Box plots representing the quantification of biofilm formation over a PVC surface of strains of interest in static conditions at 30 °C. Biofilm formation values are the proportion between the OD600 of biofilms stained with crystal violet and the OD600 of the planktonic culture. E) XY graph of the data plotted in A, B, C, and D, showing the mean values of biofilm formation for each strain at each time point tested. Error bars represent the standard deviation. Means were compared using a Brown-Forsythe and Welch ANOVA test followed by a Dunett’s T3 multiple comparison test to compare directly to the mean of the WT strain or the *ΔopaR* mutant strain. Black and maroon asterisks indicate a statistical difference compared to WT and *ΔopaR*, respectively. Adjusted P value * ≤ 0.05, ** ≤ 0.01, *** ≤ 0.001, **** ≤ 0.0001.

The control strains *ΔcpsR* and *ΔcpsS* behaved as expected. The *ΔcpsR* mutant showed a severe defect in biofilm formation. The *ΔcpsS* mutant strain showed a 195% increase in biofilm formation at the 4-hour time point, with a biofilm kinetic similar to that of the WT strain. The biofilm formed by the *ΔcpsS* mutant was almost completely dispersed and/or detached after 8 hours of growth. The differences in the biofilm formation kinetics of the *ΔopaR* and the *ΔcpsS* mutant strains could suggest that these regulators repress biofilm formation through different mechanisms and/or at different stages of biofilm development.

### OpaR interacts with the regulatory region of a variety of genes involved in c-di-GMP metabolism and biofilm formation

The genome wide landscape of OpaR-DNA interactions during exponential and stationary growth phase was recently published and the data uploaded to the Gene Expression Omnibus (GEO) Datasets with the accession number GSE122479 (36). This report did not provide a complete list of genes or DNA regions occupied by OpaR. Thus, we utilized the raw data of the ChIP-seq experiment to identified DNA regions recognized by OpaR. We then focused our attention on genes associated with biofilm formation and c-di-GMP metabolism. For this purpose, we used the tool KEGG Mapper to search for the presence of mapped objects in the *V. cholerae* Biofilm formation pathway in the dataset of genes recognized by OpaR (Fig. 9). The dataset had gene products involved in motility, biofilm matrix production, adherence to surfaces and biofilm dispersion. The genes *opaR* and *aphA* were part of the dataset of genes recognized by OpaR, supporting a previous observation of their direct regulation by this protein (37). The OpaR-ChIP-seq dataset also contained the gen *luxO* (VP2099), whose product indirectly regulates OpaR accumulation (23), and the gene VP1945, whose product is an orthologue of VarA from *V. cholerae*. VarA is a response regulator that controls the abundance of the sRNA binding protein CsrA, which indirectly modulates LuxO activity (38) and directly regulates AphA expression (39). We also identified in the ChIP-seq dataset 21 genes whose products are predicted or known to be involved in c-di-GMP metabolism (Fig. 9B). From these, three are objects in the Biofilm formation pathway of *V. cholerae:* the DGC CdgK and the PDEs CdpA and RocS (13, 40, 41). The ChIP-seq dataset also contained c-di-GMP related genes previously shown to be directly regulated by OpaR (24), and the gene *tpdA*. Our data suggest that OpaR regulates *tpdA* expression mainly through the control of c-di-GMP levels, however the ChIP-seq data could support an additional regulatory mechanism that involves direct transcriptional regulation.

**Fig. 9.**
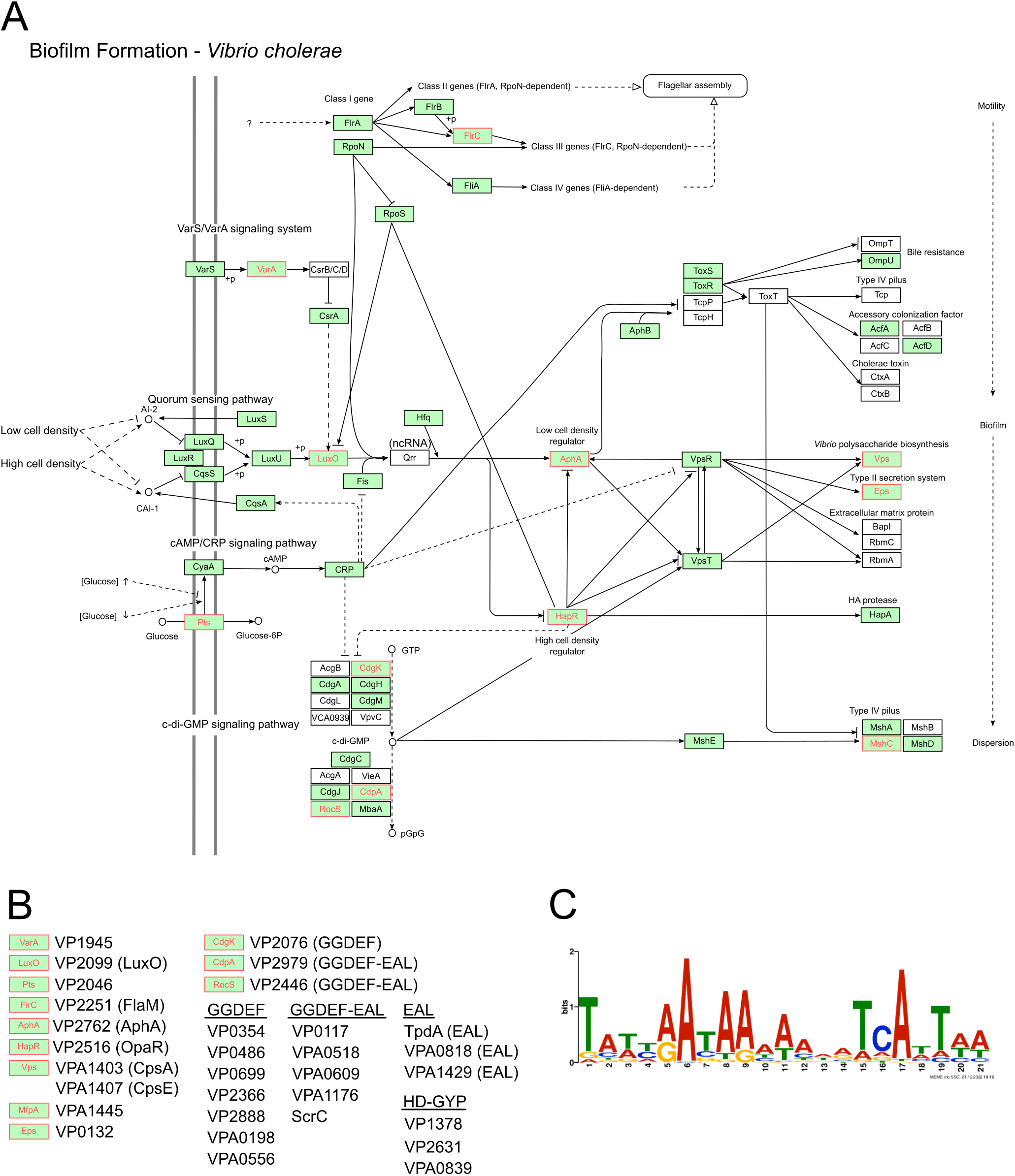
OpaR has potential regulatory roles at several branches of the biofilm regulatory circuit. A) Schematic representation of the Biofilm formation pathway generated with KEGG Mapper to illustrate gene products that are conserved in the *V. parahaemolyticus* RIMD2210633 and genes that were retrieved from a previously reported ChIP-seq experiment focused on elucidating OpaR-DNA interactions during the late-exponential and stationary growth phases (GEO accession: GSE122479) (REF PMID: 33193123). The notations are as described in the documentation for KEGG Pathways Map. Green boxes correspond to gene products that, based on the KEGG database, are also present in *V. parahaemolyticus*. Gene products labeled in red were retrieved from the analysis of the raw data of the ChIP-seq experiment mentioned above. Filled arrows indicate positive regulation or interaction while T connectors indicate negative regulation or inhibition. Dashed lines indicate indirect interaction or an uncharacterized association. Unfilled arrows indicate a connection with another Pathway map from the KEGG database. B) Biofilm and c-di-GMP-metabolism related gene products from *V. parahaemolyticus* obtained from the analysis of the ChIP-seq data for OpaR-DNA interactions. C) Logo of the conserved Motif identified within the peaks obtained from the ChIP-seq data for OpaR-DNA interactions associated to biofilm and c-di-GMP related genes. The Multiple Em for Motif Elicitation (MEME) program was used to generate the Logo.

We were able to identify a conserved OpaR binding sequence (Fig. 9C), like the one reported in the original work that generated the ChIP-seq dataset (36), in all the genes mapped to the Biofilm formation pathway of *V. cholerae* and the c-di-GMP related genes described in Figure 9B. The direct involvement of OpaR in the transcriptional regulation of *tpdA* and the multiple c-di-GMP related genes found within the ChIP-seq database deserves further attention and could give us a bigger picture of the regulatory circuitry that can be exploited by modulating OpaR abundance.

## Discussion

C-di-GMP accumulation in *V. parahaemolyticus* plays a key role in the adaptation of a surface-adhered lifestyle and surface colonization not only during biofilm formation but also during the process of swarming motility, which involves the movement over solid or semisolid surfaces using lateral flagella (27, 28, 42). Our knowledge of the mechanisms that govern changes in c-di-GMP accumulation in *V. parahaemolyticus* is limited when compared to what has been reported for *V. cholerae* (11). Nonetheless, several PDEs present in *V. parahaemolyticus* but absent in *V. cholerae* have been reported to play crucial roles in surface sensing, biofilm formation and swarming motility (25, 27–29). ScrC is the best characterized of these PDEs; in fact, ScrC is a dual-function enzyme capable of c-di-GMP synthesis and degradation. The switch in activity of ScrC is regulated through its interaction with ScrB, a periplasmic protein capable of binding a quorum-like autoinducer molecule named Signal S, produced by ScrA. ScrA, ScrB and ScrC are encoded in an operon located in the small chromosome of *V. parahaemolyticus* (VPA1513/*scrA*-VPA1512/*scrB*-VPA1511/*scrC*). The expression of this operon is crucial for the process of swarming motility and is negatively regulated by OpaR (23, 27). The PDE ScrG has also the capability to regulate cellular behaviors on solid or semisolid media, and its expression has been reported to be positively regulated by OpaR (24, 29). Lastly, the trigger phosphodiesterase TpdA, which was one of the main subjects of study in this work, was previously shown to positively modulate swimming motility and, when overproduced, to inhibit biofilm formation and promote swarming motility (25). In that work we reported that the activity of the P*_tpdA_* promoter is induced by a decrease in c-di-GMP levels but the magnitude of change in c-di-GMP that is necessary for this induction was not investigated. Although the three above mentioned PDEs affect phenotypes related to surface adaptation, little was known about the subtleties of their roles in controlling the c-di-GMP global pool. The OpaR regulated PDEs ScrC, ScrG and VP0117 showed different degrees of contribution to the upregulation of *tpdA* in the absence of OpaR. Although VP0117 was more influential than ScrG in controlling the expression of *tpdA* in the absence of *opaR*, its contribution was minor compared to the regulation exerted by ScrC. VP0117 is a GGDEF-EAL-domain containing protein whose predominant enzymatic activity has not been characterized yet, but based in these results it would appear to act as a PDE. We selected VP0117 as a potentially influential PDE in the control of c-di-GMP degradation by OpaR because it was reported to be one of the most highly expressed and induced c-di-GMP metabolizing enzymes in the absence of OpaR (24). Interestingly, the regulatory role of ScrC, ScrG and VP0117 over *tpdA* in the absence of OpaR was not accompanied by significant changes in c-di-GMP accumulation. This would suggest that the activity of the P*_tpdA_* promoter can respond to the absence of particular PDEs without major changes in the global c-di-GMP pool. We also show that the contribution of TpdA and ScrC to the control of c-di-GMP homeostasis depends on the conditions of growth; in shaking cultures, TpdA plays a major role compared to ScrC, while in cells growing over solid media the roles are swapped at the early stages of growth, but TpdA regains its dominant influence later.

OpaR has been previously shown to positively control c-di-GMP accumulation, a role that fits well with its ability to positively control CPS production and negatively control swarming motility (19, 23). However, it has also been shown that the absence of *opaR* inhibits biofilm dispersion and/or dettachment in strain BB22, and promotes biofilm formation in strain RIMD2210633 (22, 24), outcomes that would be typically associated with an elevation in c-di-GMP levels (8). These antecedents suggest that OpaR is capable of both positive and negative regulation of c-di-GMP accumulation, *cpsA* expression and biofilm formation. Is this dichotomy due to strain differences or a response to yet poorly understood environmental factors that can favor one OpaR regulatory role over the other? Although we cannot rule out differences arising from strain variability, our data strongly suggest that OpaR has this yin and yang regulatory nature in the strain RIMD2210633. Which side of its double edge sword OpaR uses to control c-di-GMP metabolism and *cpsA* expression likely depends on growth conditions such as planktonic versus surface attached, low or high population density and/or nutrient availability or the accumulation of metabolic byproducts at later stages of growth. Our data also positions TpdA and ScrC as important modulators of c-di-GMP metabolism within the regulatory circuitry of OpaR. Besides being regulated by OpaR, other temporal and functional sequestration strategies appear to control their influence in maintaining c-di-GMP homeostasis in cells growing over solid media. Temporal and functional sequestration modes of control for c-di-GMP modules refer to strategies that regulate the time for production of c-di-GMP metabolizing enzymes or associated effectors, or mechanisms that control the enzymatic activity of PDEs and DGCs allosterically (10). A result that caught our attention was that, although c-di-GMP levels were drastically reduced after 24 hours of growth over solid media in cells that lack *opaR*, the expression of *tpdA* was not induced. In contrast, in planktonic cultures a smaller decrease in c-di-GMP levels in cells lacking *opaR* was enough to induce the activity of the P*_tpdA_* promoter, even in shorter periods. This would suggest that other regulators besides OpaR and TpdA might control the expression of the P*_tpdA_* promoter in surface attached cells. The identity of these regulators and their mechanism of action will be explored in the future.

In *V. cholerae* HapR, the orthologue of OpaR, has been shown to interact and regulate a great variety of c-di-GMP metabolizing enzymes which includes both PDEs and DGCs (16, 21). So far, the absence of HapR has only been associated with an elevation in c-di-GMP levels (13, 16, 18). This would suggest that it negatively regulates the abundance of DGCs and positively regulates the abundance of PDEs, although this does not seem to be as straightforward (16, 21, 43). Based on the *in-silico* analysis of data obtained from a previously reported ChIP-seq experiment focused on OpaR-DNA interactions (36), we identified multiple DGCs, PDEs and potential dual function GGDEF-EAL proteins that could be subject of regulation by OpaR. Several of these regulatory relationships have already been explored. At least two potential DGCs, VPA0198 and VP0699, have been shown to be negatively regulated by OpaR. If in fact these proteins have DGC activity, its upregulation in the absence of *opaR* could explain why c-di-GMP levels can be increased in a *ΔopaR* mutant strain. There are at least 5 additional proteins that harbor a GGDEF domain that could be regulated by OpaR. This regulatory relationship and their enzymatic activity require further investigation to understand if all or some of them contribute to the control of c-di-GMP homeostasis by OpaR. Beside potential targets involved in c-di-GMP metabolism, targets involved in polar flagellum assembly and surface attachment such as *flaM* and *mshC* could begin to explain the biofilm dispersion phenotype associated with the absence of OpaR. Based on our results we propose that functional genomic studies of the role of OpaR on c-di-GMP synthesis and degradation and multicellular behaviors need to be designed bearing in mind the dynamic nature of the OpaR-mediated regulation of these processes.

## Material and Methods

### Strains, plasmids, and growth conditions

The strains and plasmids used in this study are described in Table 1. All strains were grown in Lysogeny Broth (LB) (1% Tryptone, 0.5% Yeast extract and 1% NaCl) or LB-agar (LB with 1.5% bacteriological agar). Strains derived from *Vibrio parahaemolyticus* RIMD2210633 were grown at 30 °C, and *Escherichia coli* strains were grown at 37 °C. Strains were grown on solid media under static conditions and in LB broth under shaking conditions at 200 revolutions per minute (rpm), at the above-mentioned temperatures. Strains and plasmids were selected by adding antibiotics to the growth media at the following concentrations: streptomycin (Str) at 200 μg/ml, kanamycin (Kan) at 30 μg/ml, gentamycin (Gen) at 15 μg/ml, chloramphenicol (Chl) at 20 μg/ml for *E. coli* and at 5 μg/ml for *V. parahaemolyticus*.

**Table 1.**
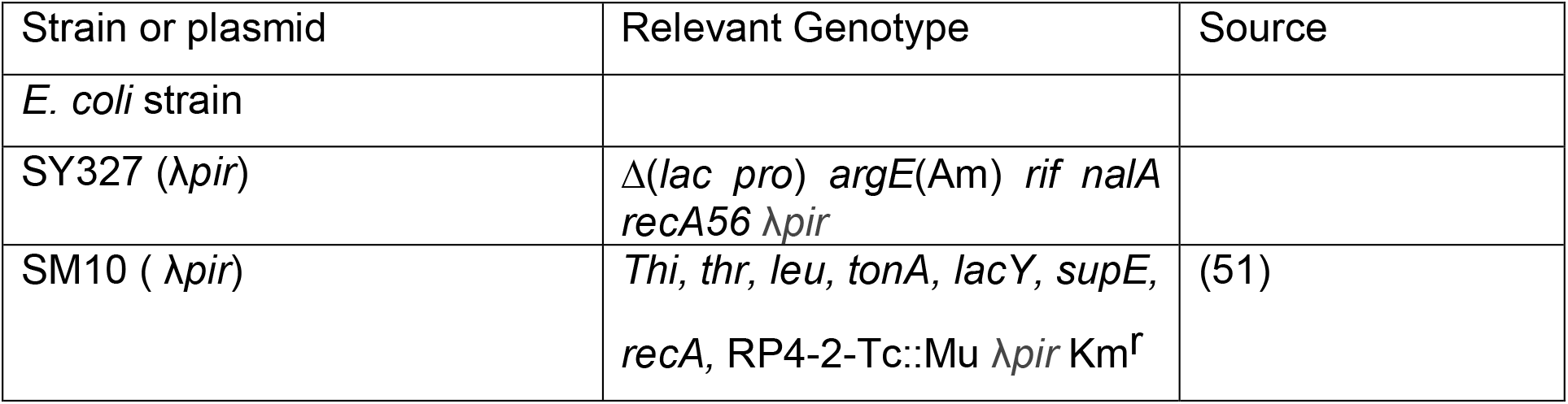
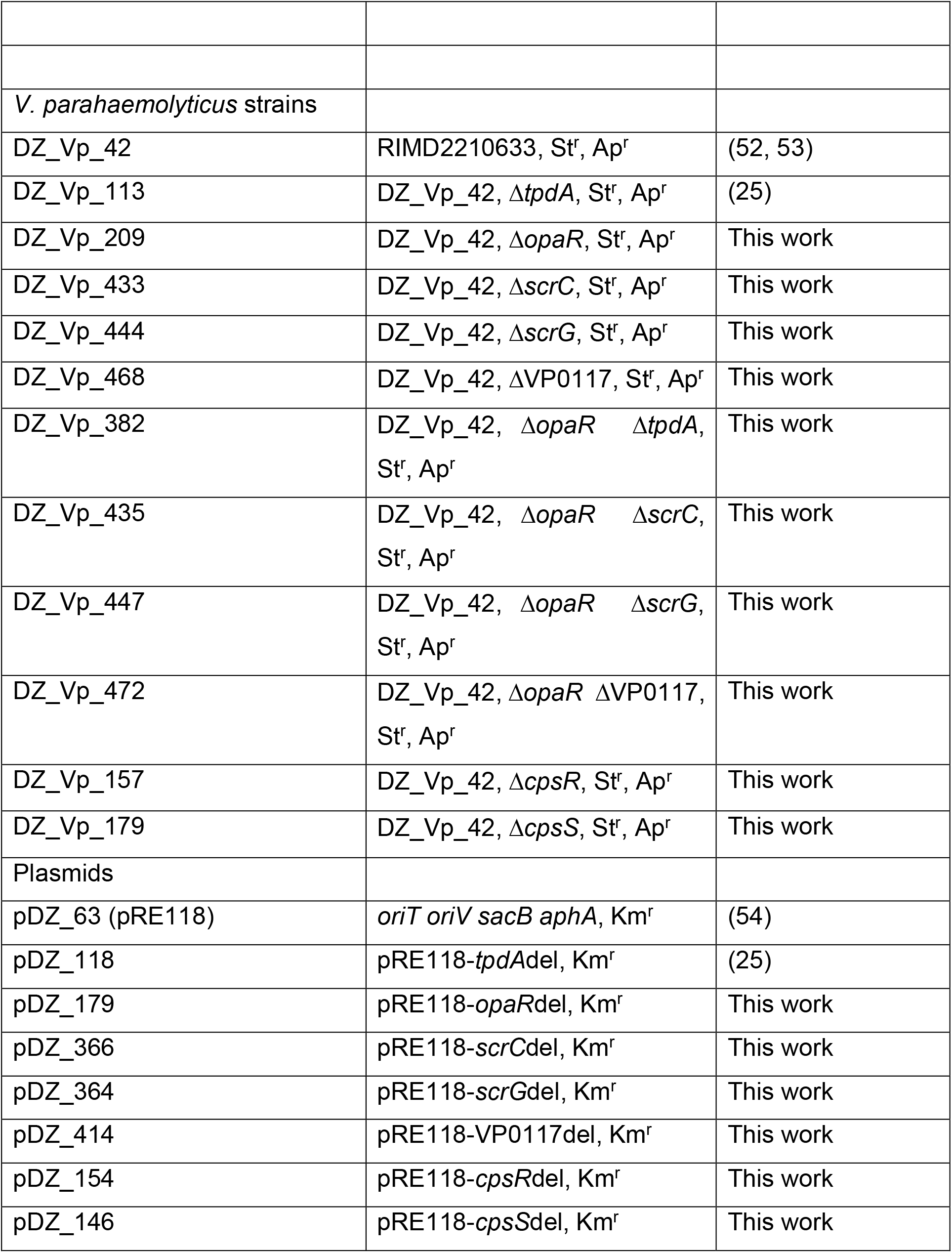
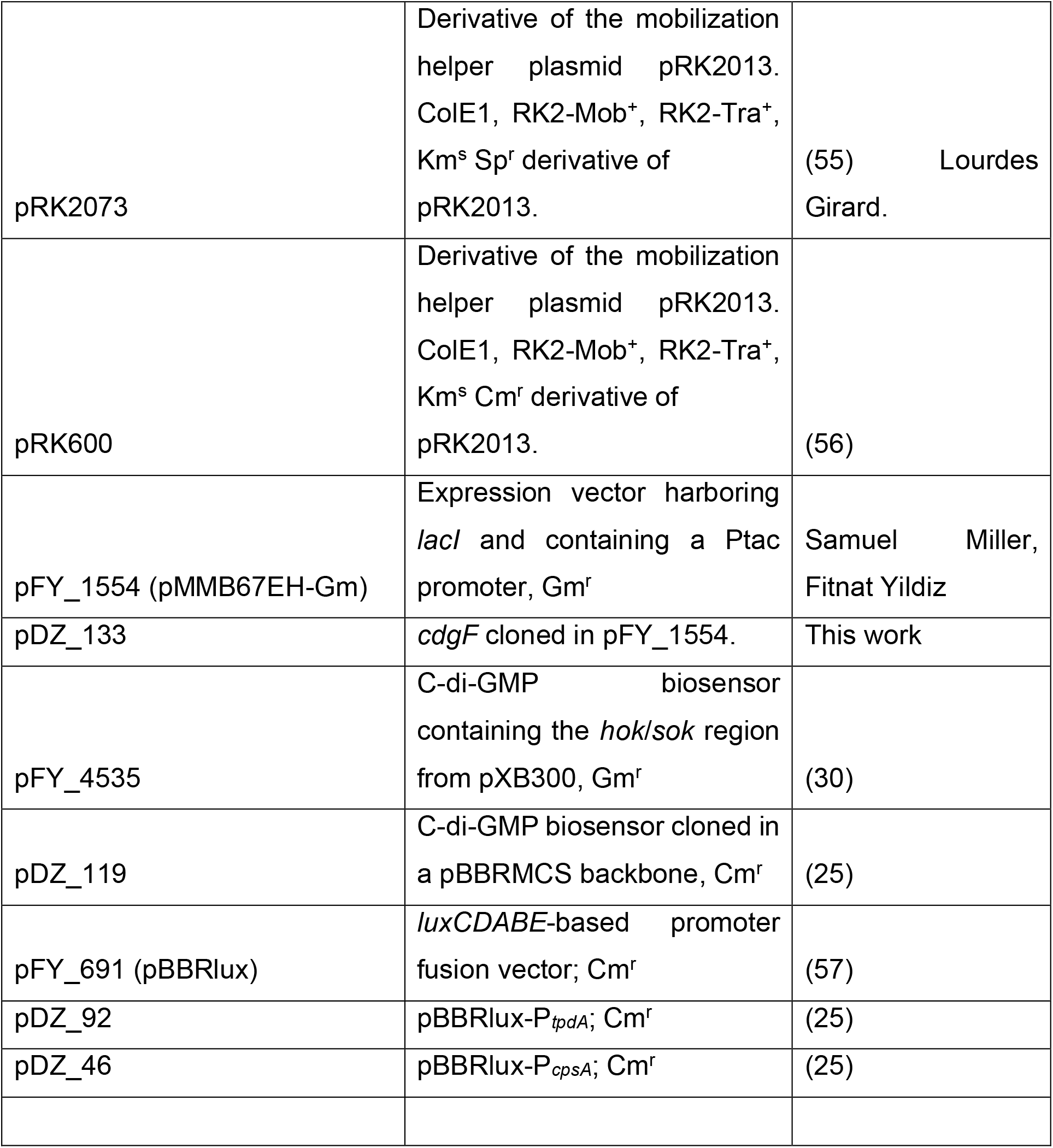
List of strains and plasmids used in this study

### Generation of genetic constructs

The primers used for the generation of genetic constructs are described in Table 2. Genomic DNA from *V. parahaemolyticus* RIMD2210633 or *V. cholerae* A1552 was used as template for the amplification of the regions of interest through PCR. PCR amplicons used for the generation of genetic constructs were produced by the high-fidelity DNA polymerase Q5 from New England BioLabs. PCR purification and plasmid isolation was done using the DNA Clean & Concentrator-5 and Zyppy Plasmid Miniprep Kits from Zymo Research, respectively.

**Table 2.**
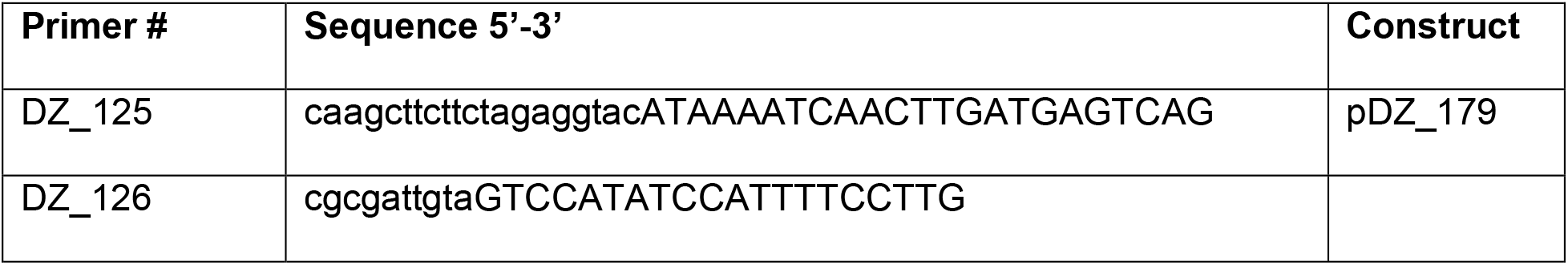
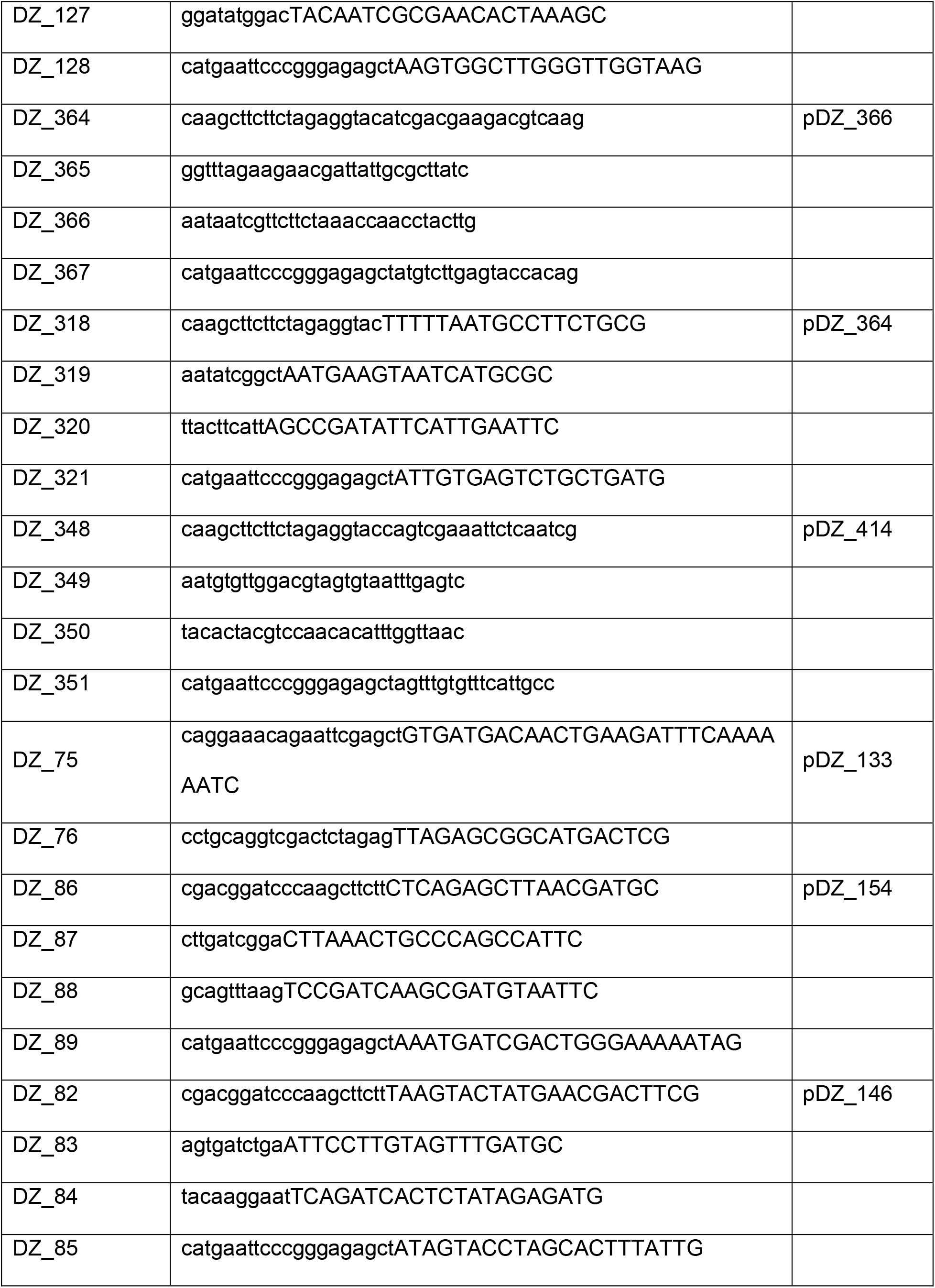
List of primers used in this study

The deletion constructs were assembled using the plasmid pRE118, which is a suicide plasmid in *V. parahaemolyticus*. The recombination substrates, used to eliminate endogenous genes through double homologous recombination, consisted of approximately 500 base pairs (bp) of upstream and downstream sequence flanking the site to be deleted. The plasmids were designed to generate in frame deletions that eliminate more than 70% of the protein sequence of the product. Plasmid pRE118 was digested with the restriction enzymes SacI-HF and KpnI-HF. The upstream and downstream recombination substrates were fused with each other and to the linearized plasmid pRE118 through a modified isothermal assembly protocol, with the NEBuilder HiFi Assembly Master Mix. We prepared 6 μl reactions consisting of at least 50 ng of each DNA assembly part and 2 μl of NEBuilder HiFi Assembly Master Mix. The primers were designed with the assistance of the online NEBuilder Assembly Tool. The assembled constructs were selected after screening clones through colony PCR with the MyTaq Red polymerase mix from Meridian Bioscience.

For the overexpression of the DGC CdgF (VCA0956) from *V. cholerae*, we cloned a PCR product corresponding to the coding sequence of *cdgF* in the plasmid pMMB67EH-Gm. The plasmid was digested with the restriction enzymes SacI-HF and BamHI-HF. The PCR product was assembled into the linearized plasmid using the same protocol of isothermal assembly described in the previous paragraph. Correct assemblies were identified through colony PCR as described above.

The genetic constructs were mobilized into the strains of interest through conjugation. For biparental mating we used the donor strain SM10λ*pir*, which carries the conjugation machinery of the RP4 conjugative plasmid. For triparental mattings we used a helper strain that host the plasmid pRK2073 or plasmid pRK600 which are derivatives of pRK2013 (44) and have the RK2 delivery machinery. To promote conjugation, we mixed the strains in a 1:1 proportion in LB broth, spotted 50 μl of the suspension over LB-agar plates without antibiotics and incubated them overnight at 37 °C. The mating spots were recovered with a sterile pipette tip, resuspended in 1 ml of LB broth and serially diluted.

The dilutions were plated over LB-agar plates with antibiotics that select for the presence of the desired plasmid in the recipient strain.

### Generation of a mutant strains with deletions of genes of interest

The deletion of the genes of interest was performed as previously described (25), with minor modifications described below. Single colonies from single recombinant strains were grown in LB broth for approximately 8 hours at 37°C with agitation (200 rpm). The cultures were then diluted (1:100) in LB broth and spread over LB-agar plates supplemented with 10% sucrose. The rest of the genetic protocol was performed as previously reported (25).

### Luminescence Assay

The luminescence assays in planktonic cultures were performed as previous described (25). To quantify light production in cells growing over LB-agar we poured 2 ml per well of LB-agar with 5 μg/ml Chl in 12-well microplates and let them dry for 1 hour with the lid open in sterile conditions, and for another 24 hours with the lid closed at room temperature. Overnight cultures of the strains of interest were diluted (1:200) in fresh LB and 2 μl of each sample was put in the middle of each well. Plates were incubated at 30°C for 24, 48, 72 and 96 hours. Luminescence of each well was measured using the Synergy H1 plate reader (BioTek). To normalize light production by cell growth, we resuspended the spot-colonies of each well in 1 ml of fresh LB. The suspension was diluted (1:5) and the optical density at 600 nm (OD600) was measured with the Epoch 2 microplate reader (BioTek). The Relative Luminescent Units (RLU) are calculated as arbitrary luminescence divided by the OD600 of the resuspended spot-colonies. Two independent experiments were done with three biological replicates each.

To analyze luminescence from biofilms formed over glass tubes under shaking conditions we first prepared overnight cultures of the strains of interest. The test tubes containing bacterial cultures were always set angled in a tube-rack inside a shaking incubator. The overnight cultures were diluted (1:1000) in 5 ml of LB broth with 5 μg/ml Chl and grown with agitation at 30 °C for 24 hours. Under these conditions, biofilms are formed a few centimeters above the planktonic culture. A sample of the planktonic culture was taken and diluted (1:5) to measure the OD600 as described above. The rest of the planktonic culture was carefully discarded avoiding disrupting the biofilms. The test tubes with the biofilms were washed with 1 ml of LB to remove the remaining planktonic culture. Biofilms were resuspended in 800 μl of LB. Two hundred μl samples of each resuspended biofilm were transferred to white, 96-well plates with clear bottom. We measured the OD600 of the resuspended biofilms and their light emission using the Synergy H1 plate reader (BioTek). The RLU are calculated as arbitrary luminescence units per ml divided by OD600 of the biofilm samples. Experiments were performed twice independently with three biological replicates each time.

### Determination of relative abundance of c-di-GMP using a genetic c-di-GMP reporter

The estimation of c-di-GMP levels in planktonic cultures using the c-di-GMP genetic reporter present in plasmids pFY4535 and pDZ119 was performed as described previously (25). To measure c-di-GMP levels using this reporter in cells grown over LB-agar, we performed the same type of experiments as described above for the luminescence assays with minor modifications. Overnight cultures of the strains of interest were diluted (1:200) in fresh LB and 2 μl of each sample were pipetted in the center of wells containing LB-agar and 15 μg/ml Gen. Plates were incubated at 30 °C for 24, 48, 72, 96 and 120 hours. Fluorescence of the Amcyan and Turborfp proteins produced by the spot colonies from each well was measured using the Synergy H1 plate reader (BioTek). We used the constitutive fluorescence of Amcyan to normalize TurboRFP production by reporter expression, hence we were able to track fluorescence from the same spot colonies over time. The Relative Fluorescence Intensity (RFI) was calculated by dividing the arbitrary units of fluorescence intensity of TurboRFP by the arbitrary units of fluorescence intensity of Amcyan. Three independent experiments were done with a total of at least 5 biological replicates. To analyze c-di-GMP levels from biofilms formed over glass tubes under shaking conditions using the c-di-GMP genetic reporter we followed a similar experimental procedure as the one described for the luminescence assays. We first grew overnight cultures of the strains of interest, diluted them (1:1000) in 5 ml of LB broth with 15 μg/ml Gen and grew them again with agitation at 30 °C for 24 hours. The planktonic culture was carefully discarded avoiding disrupting the biofilms. We washed the biofilms with 1 ml of sterile distilled water to remove the remaining planktonic culture. We then resuspended the biofilms in 800 μl of sterile distilled water and transferred a sample of 200 μl to black, 96-well plates. We measured the fluorescence of Amcyan and TurboRFP from resuspended biofilms using the Synergy H1 plate reader (BioTek). To calculate biofilm formation units the arbitrary units of fluorescence intensity of Amcyan from the biofilms was divided by the OD600 of their corresponding diluted (1:5) planktonic cultures. Experiments were performed twice independently with three biological replicates each time.

### Liquid-Solid interface Biofilm assays over a PVC surface

Biofilm formation over Polyvinyl Chloride (PVC) wells was performed using a previously reported crystal violet staining procedure with some modifications (45). The strains of interest were grown overnight in 5 ml of LB. Cultures were then diluted to an OD600 of 0.02. One hundred and fifty μl samples of the cultures and a blank sample consisting of sterile LB broth were transferred to 96-well PVC microplates previously sterilized with UV light for 10 minutes. Wells were incubated for 2, 4, 6 and 8 hours at 30 °C under static conditions. To measure the growth of the planktonic cultures we analyzed their OD600 in the Epoch2 microplate reader. The planktonic cultures were carefully discarded in a 10% bleach solution and washed twice with tap water by decantation. The wells were dried upside down over paper towels and afterwards the biofilms were stained with a 0.1% crystal violet solution for 10 minutes at room temperature. To solubilize the dye retained by the biofilms we added 150 μl of ethanol (absolute) to the wells and incubated for approximately 30 minutes at room temperature under static conditions. We measured the OD600 of the solubilized crystal violet dye with the Epoch2 microplate reader. Biofilm formation units were obtained by dividing the OD600 of the dyed retained by the biofilms by the OD600 of the planktonic cultures from the same well. At least three independent experiments were performed for each strain of interest.

### Bioinformatic analysis of the GSE122479 experiment from the Gene Expression Omnibus (GEO) Datasets

We used the raw data of the GEO datasets with the accession number GSE122479 generated in a previous report (36). The analysis was performed by the bioinformatic facility “Unidad de Análisis Bioinformáticos” from the Center for Genomic Sciences (UNAM). The data was analyzed with FastQC v0.11.8 before trimming (https://www.bioinformatics.babraham.ac.uk/projects/fastqc/). The program trim_galore 0.6.6 (https://www.bioinformatics.babraham.ac.uk/projects/trim_galore/) was used to remove adaptor and other undesirable sequences. The obtained sequences were mapped to the reference genome (GCF_000196095.1_ASM19609v1) using the program Bowtie2 (46). The programs MACS2 and HOMER were used for peak calling.

The genes associated to the ChIP-seq peaks identified were mapped to genes in the KEGG Pathway database (https://www.genome.jp/kegg/pathway.html) using KEGG Mapper (47, 48).

The sequences of the peaks (or the reverse complementary sequence for genes codified in the negative strand) associated to genes mapped to the Biofilm formation KEGG Pathway or gene products predicted to be involved in c-di-GMP metabolism, were used as seed to identified a conserved motif using the Multiple Em for Motif Elicitation (MEME) program (49, 50).

## Acknowledgements

We are thankful to DGAPA-PAPIIT (UNAM) for the financial support provided through the project IA201821, awarded to David Zamorano-Sánchez.

We acknowledge the technical assistance provided by Eugenio Lopez-Bustos, Paul Gaytan, Santiago Becerra, and Jorge A. Yañez, from Unidad de Síntesis y Secuenciación, Instituto de Biotecnología-UNAM, in oligonucleotide synthesis and DNA sequencing. We are thankful to Mishael Sánchez-Pérez from the Unidad de Análisis Bioinformático, Centro de Ciencias Genómicas-UNAM, for his technical assistance in the *in-silico* analysis of the GEO Datasets with reference number GSE122479.

